# *Puya*: A new example of adaptive radiation in the Northern Andes

**DOI:** 10.64898/2026.11.24.768968

**Authors:** Nicolás Castillo-Rodríguez, Julián Aguirre-Santoro, Edgardo M. Ortiz, Rachel S. Jabaily, Rosana Zenil-Ferguson

**Affiliations:** Department of Biology, University of Kentucky, Lexington, KY 40508; Missouri Botanical Garden, St. Louis, MO 63110; Florida Museum of Natural History, University of Florida, Gainesville, FL 32611; Department of Organismal Biology & Ecology, Colorado College, Colorado Springs, CO 80903

**Keywords:** Competitive exclusion, diversification, character displacement, pulsed evolution, sky-islands

## Abstract

- Most documented examples of plant adaptive radiations occur on oceanic islands, but high-elevation mountain systems offer similar ecological opportunities. Here, we investigate the genus *Puya* (Bromeliaceae), a rapidly diversifying Andean lineage associated with high-elevation ecosystems, to assess how ecological opportunity and biotic interactions shape diversification and morphological evolution in fragmented mountain landscapes.
- Using a phylogenomic dataset for the Septentrional Clade of *Puya*, a clade largely restricted to the páramo habitat, we performed biogeographic reconstructions and trait evolution analyses using parametric models.
- We identified a temporal increase in diversification rates consistent with the onset of páramo expansion, together with a clade-specific increase in lineages associated with this ecosystem. Likewise, we found that character displacement and competitive exclusion drove phenotypic divergence and identified signatures of pulsed evolution.
- We propose the Septentrional Clade of *Puya* as a new example of adaptive radiation in continental sky-island systems and demonstrate that ecological opportunity and biotic interactions jointly shaped its evolutionary history. Our findings provide new insights into how ecological pressures shape macroevolutionary patterns in highly diverse plant clades and show that mountain radiations can generate morphological diversity through processes comparable to those documented in classic examples of adaptive radiations in oceanic islands.

## Introduction

Biotic interactions influence both biogeographic and phenotypic evolution during diversification through pre- and post-sympatric effects on community assembly. Some biotic interactions, like competitive exclusion, may prevent the colonization and long-term coexistence of species that share the same niche (Stroud *et al*., 2019, 2024; Quintero & Landis, 2020). Nonetheless, post-sympatric character displacement of traits can reduce interspecific competition and local extinction (Brown & Wilson, 1956; Eisen & Geber, 2018; Stroud *et al*., 2019; Morita & Yamamichi, 2023). Character displacement in sympatric communities primarily arises from two types of interactions: i) reinforcement of reproductive barriers, which prevents reproductive interference resulting from incomplete species recognition, and ii) ecological divergence, which reduces competition for shared resources among overlapping populations (Brown & Wilson, 1956). Phenotypic divergence is often correlated with the occupation of novel environments in adaptive radiations, where rapid bursts of diversification of ancestral lineages allow new species to explore a wide range of resources and ecological niches (Losos, 2010; Gillespie *et al*., 2020). These dynamics have been extensively documented in lineages that diversified on oceanic islands, with iconic examples such as *Anolis* lizards in the Caribbean (Losos *et al*., 2006) and Hawaiian silverswords (Witter & Carr, 1988). These well-studied systems have served as empirical foundations for the development and validation of probabilistic biogeographic and evolutionary models of insular lineage diversification (Sanmartín *et al*., 2008; Ronquist & Sanmartín, 2011; Landis *et al*., 2018). Although examples of adaptive radiations in the mainland exist (Seehausen, 2006; McGuire *et al*., 2014; Merrill *et al*., 2015), adaptive radiations in continental systems, and the factors that influenced them, remain comparatively understudied.

An example of ecological and geographic isolation in continental formations is high-elevation mountaintops. Commonly referred to as sky-islands, these habitats are comparable to oceanic archipelagos in terms of isolation and can promote rapid diversification (Vuilleumier, 1970; Brown, 1971, 1978; Love *et al*., 2023). In particular, the Andean páramo has long been proposed as a sky island system and potential hotspot for adaptive radiations, supporting exceptional species richness and phenotypic diversity driven by environmental adaptation (Cortés *et al*., 2018; Guevara-Andino *et al*., 2025). Its biota is characterized by high levels of plant diversity and endemism that originated through complex evolutionary processes during the last 3–5 My (van der Hammen & Cleef, 1986; Sklenář *et al*., 2014; Londoño *et al*., 2014). An excellent model lineage to study evolutionary patterns in páramo is the Septentrional Clade of *Puya* (Bromeliaceae), which began diversifying in the Northern Andes during this period and currently comprises approximately 40 species, most of them restricted to páramo ecosystems (Aguirre-Santoro *et al*., 2024). This clade exhibits wide variation in both vegetative and reproductive traits (26), making it an ideal system for evaluating the effects of ecological opportunity, biotic interactions, and rapid species diversification.

In this work, we assess whether diversification dynamics of *Puya* are associated with páramo formation and whether biotic interactions shaped the observed phenotypic divergence in the clade associated with this ecosystem. We leverage recent phylogenomic work and incorporate new genomic data to obtain an updated phylogeny and perform biogeographic and trait evolution analyses for the Septentrional Clade of *Puya*. We predict that ecological opportunity promoted rapid diversification, whereas interspecific competition contributed to phenotypic divergence among sympatric species.

## Materials and Methods

### Phylogenetic inference and divergence time estimation

We used two phylogenetic trees representing distinct analytical scales. First, we used the phylogeny published by Aguirre-Santoro *et al*. (2024) for our diversification analyses. This first phylogeny has 85 lineages, representing approximately 35% of the described species diversity of the genus, with 25 species belonging to the Septentrional Clade. For the biotic interaction analyses, focused on the Septentrional Clade, we analyzed genomic data from 38 *Puya* individuals, including 34 from the Septentrional Clade, representing 26 formally recognized species and four undescribed taxa, accounting for nearly 80–85% of the clade’s diversity. For *P. cryptantha* and *P. nitida*, which have broader geographic distributions, three populations were sampled per species. The remaining four samples of *Puya* are not part of the Septentrional Clade but were included as outgroups to root and date our phylogeny proposal. The genomic data for 26 of the samples were already used in the phylogenomic study of *Puya* published by Aguirre-Santoro *et al*. (2024); thus, DNA extraction, hybrid enrichment, and sequencing for the remaining 12 samples were performed using the methodology described in the same publication.

Sequence data were generated using the Angiosperms353 probe set (Johnson *et al*., 2019) and processed with the Captus pipeline v1.3.2 (Ortiz *et al*., 2026). Raw reads were cleaned and assembled *de novo* using the “clean” and “assemble” modules, respectively, under default parameters. Angiosperms353 loci were then retrieved via the “extract” module with default settings, and subsequently aligned using the “align” module with the flags --min_samples 2 --min_coverage 0.66 -f AA,NT,GE,GF. To improve recovery consistency and overall yield, the resulting GF (full gene and flanking regions) alignments were used as input for the “new_targets_from_alignments” module to generate *Puya*-specific targets, applying a 70% clustering threshold (-i 70). These custom targets were retrieved using the “extract” module with --nuc_min_identity 90. Finally, the newly retrieved loci were aligned using the “align” module with the settings --align_method mafft_genafpair --clipkit_gaps 0.50 --min_coverage 0.50.

Individual GF locus trees were inferred using maximum likelihood in IQ-TREE v2.0.6 (Minh *et al*., 2020). For each locus, model selection was performed using the -m TEST flag based on the Akaike Information Criterion (AIC), and branch support was assessed with 1000 ultrafast bootstrap replicates (-bb 1000). To account for incomplete lineage sorting, a species tree was estimated from the resulting gene trees in a coalescent framework using ASTRAL-Pro v1.1 (Zhang *et al*., 2020). Robustness of the species tree topology was evaluated using local posterior probabilities, and quartet statistics were retained for downstream analysis.

We obtained a time-calibrated tree by first selecting a subset of 25 GF loci using SortaDate (Smith *et al*., 2018) to reduce the dataset’s complexity. We performed the divergence-time estimation in BEAST 2.7.7 (Bouckaert *et al*., 2019) using the previously inferred species tree as a fixed topology. We applied the substitution models identified with ModelFinder (Kalyaanamoorthy *et al*., 2017) for each of the 25 GF loci, kept the tree topologies linked while allowing substitution rates to vary independently, and assumed a log-normal relaxed molecular clock. We incorporated two normally distributed secondary calibration priors: 10.1 Ma for the crown age of the Puyoideae and 2.8 Ma for the crown age of the Septentrional Clade of *Puya*. These priors are derived from Aguirre-Santoro et al. (2024) and Givnish *et al*. (2011), the latter including fossil calibrations across all monocots. The analysis was conducted using two independent Markov Chain Monte Carlo (MCMC) runs of 50 million generations with sampling every 100,000 generations and a burn-in fraction of 25%. Finally, we produced a maximum clade credibility (MCC) tree.

### Diversification analyses

We implemented two distinct approaches to estimate diversification rates across the phylogeny of *Puya*. First, we used an EBD (Stadler, 2011; Höhna, 2015) model in RevBayes (Höhna *et al*., 2016) to identify diversification-rate variation through time. The EBD model partitions time into intervals, each with its own constant speciation and extinction rates. Following Magee *et al*. (2020), we applied the horseshoe Markov random field as a non-parametric prior on the log-transformed diversification rates. Diversification rates at the present were defined following the method of moments estimate (Magallon & Sanderson, 2001), in which the expectation of the speciation rate at the crown-group age under a pure-birth model equals 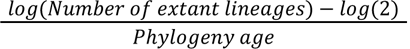. We obtained speciation and extinction rates for each interval by exponentiating the sum of the present diversification rate and the cumulative log-rate changes across intervals. We defined a total of 25 evenly spaced intervals and included a sampling fraction representing the 38% taxon sampling of the genus. To check for convergence, two MCMC chains were estimated for 40,000 generations, at which point all parameters in both chains reached effective sample sizes (ESS) greater than or equal to 250. Likewise, we performed a ClaDS analysis (Maliet *et al*., 2019; Maliet & Morlon, 2022) using the PANDA package available in Julia (Bezanson *et al*., 2017) to test whether the Septentrional Clade has higher branch-specific diversification rates on *Puya*. This method assumes that each lineage has a specific diversification rate by modeling a birth-death process in which, at cladogenetic events, diversification rates are inherited with a shift. We used the same sampling fraction as in the EBD and the prior distributions described by Maliet & Morlon (2022).

### Biogeographic analyses

Reconstructing the biogeographic history of the Septentrional Clade of *Puya* is key to determining the timing of sympatry and colonization events. We defined nine areas by considering the páramo bioregions proposed by Londoño *et al*. (2014) according to parsimony analyses of endemicity in Colombia, followed by subdividing some of these bioregions based on *Puya* endemism. We then accounted for the geological history of each region, where deep valleys prevented páramo connectivity even during glacial periods (van der Hammen, 1974; Hooghiemstra *et al*., 2006; Flantua *et al*., 2019), and included the páramo areas in the Cordillera de Mérida, Venezuela, and the high-cloud ecosystem in the Cordillera de Talamanca, Costa Rica. Using specimen records from the Global Biodiversity Information Facility, JSTOR Global Plants (https://plants.jstor.org/), and the herbaria B, COL, CUVC, F, FMB, HECASA, HUA, MA, MO, K, NY, P, and WU, we assigned *Puya* species to one or more of the biogeographic regions that were considered. Later, a DEC+J model (Matzke, 2014) was implemented in RevBayes (Höhna *et al*., 2016) after following a model selection approach using the minimum corrected Akaike Information Criterion (AICc) in BioGeoBEARS (Matzke, 2013, 2014) (Supporting Information, Table S3). We set the highest number of areas that can be occupied at a given time by a lineage to three, as our extant species indicate, and defined the root state frequencies to a value greater than zero for those states containing the southern regions in the assessed distribution (the Central Cordillera and its combinations that include the Eastern Cordillera and Farallones de Cali), following Aguirre-Santoro et al. (2024).

### Morphological traits description for sympatric species

We modeled phenotypic variation in the Septentrional Clade of *Puya* to understand how morphological change occurred. The input for phenotypic models was vegetative and reproductive traits. The vegetative traits (Fig. 1) were: i) plant length (mm), defined as the distance from the rosette base to the inflorescence apex; ii) leaf blade area (mm²), calculated as the average of all possible combinations of measured leaf blade lengths and leaf basal width, assuming a triangular shape characteristic of *Puya* leaves. Additionally, the reproductive traits were: i) total number of flowers per inflorescence; ii) floral bract ratio, described as the mean floral bract length relative to mean floral basal width; and iii) sepal ratio, expressed as the mean sepal length relative to mean sepal basal width. Morphological measurements were obtained from species descriptions or directly from the examined specimens across herbaria. These traits were measured in one to five individuals per species and averaged for downstream analyses to account for intraspecific variation.

**Figure 1.**
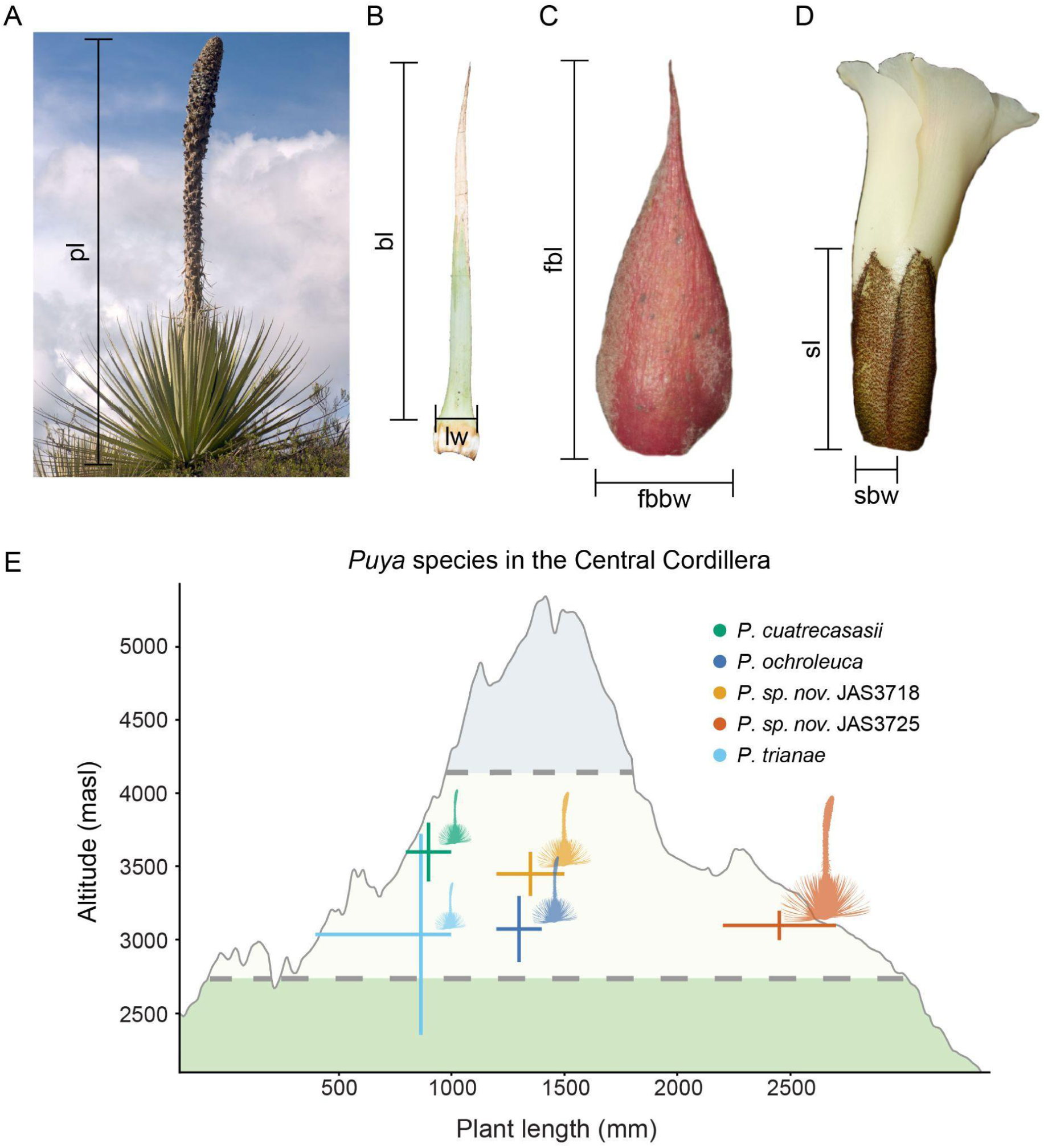
Morphological traits evaluated in *Puya*. (A) Plant length (PL). (B) Leaf blade area, measured from blade length (BL) and leaf width (LW). (C) Floral bract ratio, calculated from floral bract length (FBL) and floral bract basal width (FBBW). (D) Sepal ratio, calculated from sepal length (SL) and sepal basal width (SBW). (E) Relationship between plant length and elevation for *Puya* species inhabiting the Central Cordillera. Points represent mean trait values, and horizontal lines indicate the observed range for each species.

In addition, we described whether sympatric species differ in their altitudinal distribution for each morphological trait, as an indicator of potential ecological differentiation. For this analysis, we considered the nine previously defined areas and additionally included nine species that were not sampled in the phylogenetic reconstruction, since this comparison does not rely on the phylogeny.

### Evaluating the effects of sympatry

We used the TRIBE model implemented in Julia (Bezanson *et al*., 2017) through the package Tapestree (Quintero & Landis, 2020). This model, developed within a Bayesian framework, simultaneously accounts for the interdependence of trait evolution and biogeography. It assumes sympatry of daughter lineages that inherit the same ranges occupied by their immediate ancestor, but it does not account for allopatric cladogenesis that may have happened in the páramos. The TRIBE model estimated three key parameters that assess the effect of biotic interactions on character change (ω_x_), colonization (ω_1_), and local extirpation rates (ω_0_), and allowed us to infer processes of character displacement, competitive exclusion, or environmental filtering against a null case scenario where biotic interactions do not have any effect. To check for parameter convergence and an ESS for all parameters higher than 250, we ran two MCMC chains for each trait in our dataset for 300,000 generations.

### Vegetative and reproductive trait evolution

Understanding how our vegetative and reproductive traits have evolved through time requires testing different models of evolution for continuous traits. We used the R (R Core Team, 2024) package pulsR (Landis & Schraiber, 2017) to fit Gaussian models for continuous trait evolution (Brownian Motion (BM), Ornstein-Uhlenbeck (OU), and Early Burst (EB)), non-Gaussian models that describe semi-continuous trait evolution but include the possibility of sudden phenotypic changes (Jump Normal (JN), Normal Inverse Gaussian (NIG), and Variance Gamma (VG)), and models that include a component of BM diffusion to these pure-jump models (BM+JN, BM+NIG, and BM+VG). We computed the AICc and its corresponding weight (wAICc) to identify the model that best explains the evolution of each *Puya* trait. We chose the best model for each trait if the best wAICc score of a given model was at least twice as high as that of the rest. However, as noted by Landis & Schraiber (2017), this criterion may lack sensitivity when comparing a large number of models. Consequently, among Lévy process models, only the one with the highest AICc was selected for representation. To ensure that true maximum-likelihood estimates were obtained, we conducted 10 independent parameter searches for each trait analyzed. Once a model was selected, we used the chosen model to simulate the stochastic mapping of each trait using the package creepy-jerk (Landis *et al*., 2013) and show where Lévy processes were supported. The software creepy-jerk models Lévy processes that include a BM diffusion without including BM+NIG; consequently, for this last particular model, it was approximated to BM+JN and BM+VG. We ran the stochastic mapping simulations twice for each trait to check for parameter convergence and for 500,000,000 generations. We deleted the first 25% of each run corresponding to the burn-in stage. On the other hand, if a trait was better adjusted to a Gaussian model, the stochastic maps were calculated using the contMap function in the R (R Core Team, 2024) package phytools (Revell, 2012) since it generates estimates that correspond to the expectation under a BM process.

## Results

### Diversification analyses

We found temporal and branch-specific heterogeneity in diversification rates across *Puya* that may reflect the same macroevolutionary signal associated with the Septentrional Clade and the páramo formation (Fig. 2). An Episodic Birth–Death (EBD) (Stadler, 2011; Höhna, 2015) analysis indicates that while relative extinction rate estimates remain constant through time, net diversification rates increased between 6–3 Ma, followed by a decline toward the present. Consistently, branch-specific diversification estimates from a cladogenetic rate shift analysis (ClaDS) (Maliet *et al*., 2019; Maliet & Morlon, 2022) reveal elevated rates in Northern Andean lineages, particularly within the Septentrional Clade, whose radiation coincides with that rate peak. Diversity and lineage-through-time plots further support this diversification increase. Together, these patterns are consistent with a diversification shift driven by the ecological opportunity offered by páramo formation in the Northern Andes during the late Miocene and Pliocene (van der Hammen & Cleef, 1986; Sklenář *et al*., 2014; Londoño *et al*., 2014), followed by progressive niche filling.

**Figure 2.**
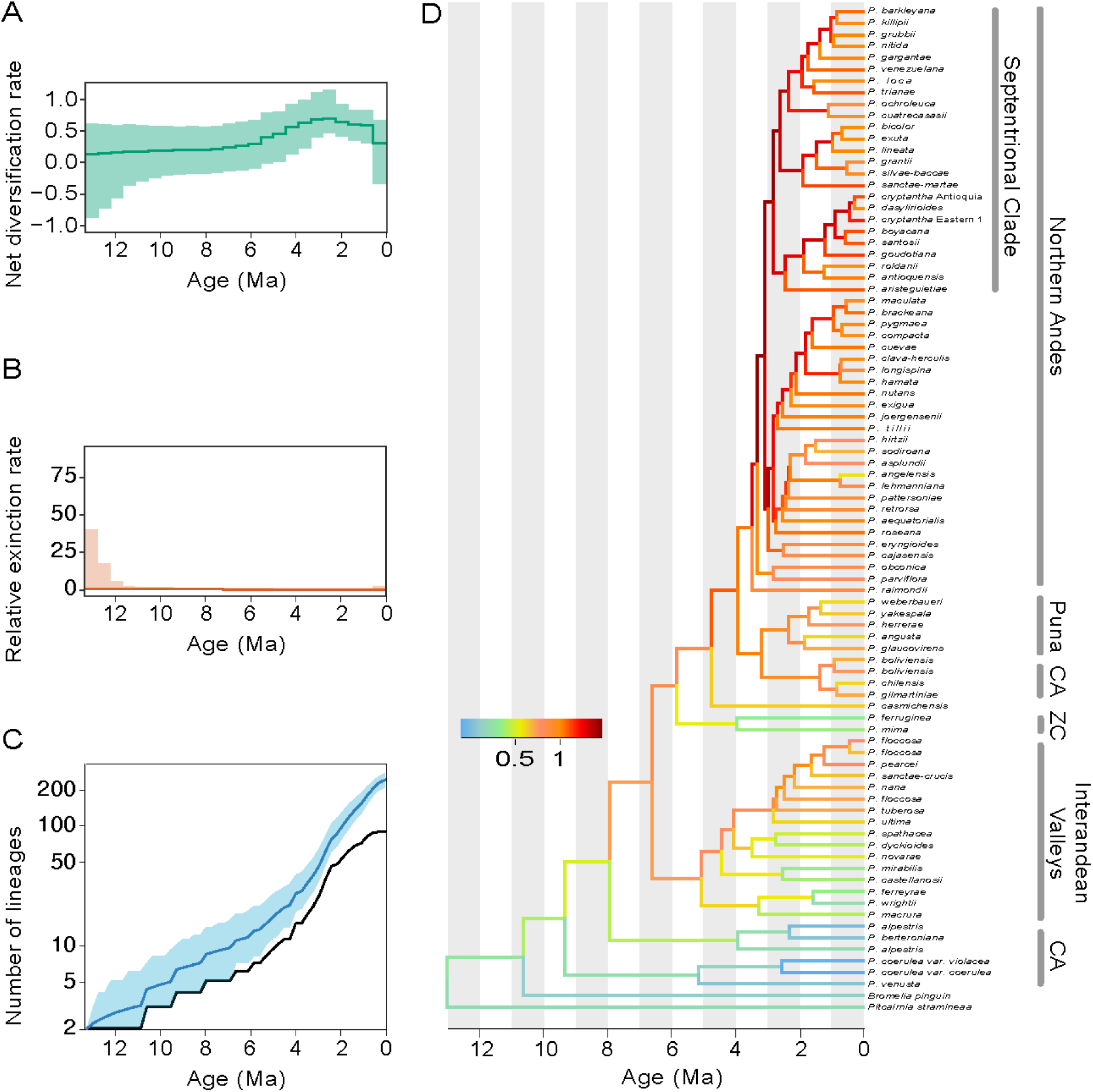
Diversification estimation in *Puya*. (A and B) Net diversification and relative extinction rates estimated under the EBD model. Thick lines indicate posterior median estimates, and shaded areas represent 95% credible intervals. (C) Diversity through time plot from the ClaDS analysis. The maximum a posteriori estimate is shown in blue, with the 95% credible interval shaded in light blue. The lineage-through-time curve for the observed phylogeny is shown in black. (D) Time-calibrated phylogeny of *Puya*, with branches colored according to lineage-specific diversification rates estimated with ClaDS. CA indicates Central Andean species, and ZC indicates the zygomorphic clade.

### Phylogenetic reconstruction and biogeographic analyses

We updated a phylogenetic tree for the Septentrional Clade and recovered relationships consistent with previous studies but with significantly improved internal resolution and support that resulted from expanded sampling and increased DNA sequence recovery using the Angiosperm353 (Johnson *et al*., 2019) dataset with the Captus (Ortiz *et al*., 2026) software. Despite improved resolution, quartet support revealed substantial gene discordance across all nodes, and widespread, multi-accessioned species such as *Puya cryptantha* and *P. nitida* were recovered as non-monophyletic (SI Appendix, Fig. S1). The time-calibrated phylogeny describes how the Septentrional Clade of *Puya* radiation started 3.7 Ma (95% Highest Probability Density (HPD): [2.7-4.7]; SI Appendix, Fig. S2), which corresponds to the formation of páramo 3–5 Ma, when Andes elevations reached a level suitable for the ecosystem (van der Hammen & Cleef, 1986; Sklenář *et al*., 2014; Londoño *et al*., 2014). Ancestral state estimation under the Dispersal, Extinction, and Cladogenesis model with jumps (DEC+J) (Matzke, 2014) supports a distribution spanning the Central Cordillera and the Eastern Cordillera at the root of the tree (Fig. 3). Early nodes in the phylogenetic tree support geographic subdivision between these mountain systems, followed by repeated dispersal events into isolated páramo regions between 1.5 and 3 Ma. Within the last million years, subsequent dispersal events from the Eastern Cordillera occurred, including the arrival of *Puya* to the Cordillera de Talamanca, the most isolated region currently occupied by the genus. This intricate process of *Puya* diversification illustrates a recent pattern of multiple colonizations in the páramo sky-islands. In addition, the cladogenetic parameters used in the DEC+J model indicate that a higher proportion of speciation events occurred in sympatry (mean: 0.754; 95% HPD: [0.395, 0.987]), followed by allopatry (mean: 0.229; 95% HPD: [0.001, 0.594]), and a small proportion as jump dispersals (mean: 0.017; 95% HPD: [0.001, 0.034]; SI Appendix, Fig. S3).

**Figure 3.**
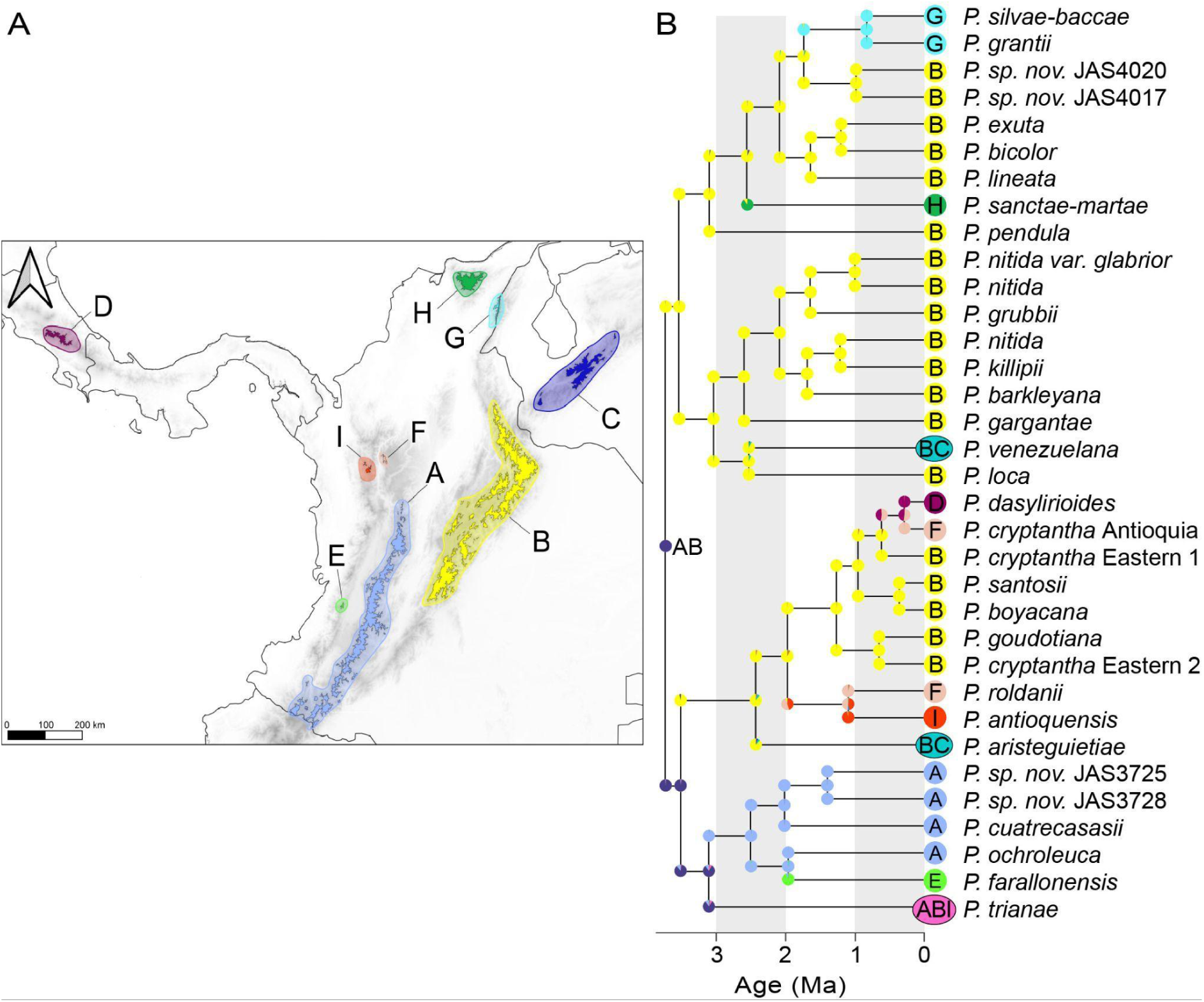
Biogeographic reconstruction for the Septentrional Clade of *Puya*. (A) Geographic areas were designated as follows: A, Central Cordillera; B, Eastern Cordillera; C, Cordillera de Mérida; D, Cordillera de Talamanca; E, Farallones de Cali; F, Páramo de Belmira; G, Serranía del Perijá; H, Sierra Nevada de Santa Marta; and I, Western Antioquia Páramo. (B) Phylogeny with ancestral range reconstruction. Tip labels indicate the geographic regions in which each species occurs, and colors in nodes and shoulders show the ancestral state estimation under the DEC+J model for the nine regions, assuming a maximum of three regions per node.

### Effects of sympatry on morphological traits

The Septentrional Clade shows distinct niche use in sympatry based on the assessed traits (Fig. 1 and SI Appendix, Figs. S4-S8). The relationship between altitude and the evaluated traits shows that sympatric species in the same altitudinal range tend to differ phenotypically. Likewise, species with overlapping or similar trait values tend to occur at different elevations. Moreover, this pattern remains consistent across most areas, suggesting that competition may shape trait divergence. In the case of the Eastern Cordillera, this mountain chain has the highest species richness and exhibits greater overlap both in altitude and trait values among lineages. Nonetheless, it has the broadest availability of ecological niches.

Moreover, using the Trait and Range Interspecific Biogeographic Evolution (TRIBE) model (Quintero & Landis, 2020), we identified two distinct patterns in which biotic interactions have shaped the evolution of the Septentrional Clade of *Puya* (Fig. 4A and B and SI Appendix, Fig. S9 and Table S1). First, we found that plant length, leaf area, and flower number diverge in sympatry, with P(ω_x_ < 0) ≥ 0.95. This pattern supports character displacement because of post-sympatric biotic interactions. These three traits did not show an effect of biotic interactions on colonization or extirpation rate parameters, as our inference did not differ from expectations under a Brownian Motion (BM) process, with the 95% HPD intervals for ω_1_ and ω_0_ including zero. In contrast, although changes in floral bract and sepal ratios did not differ from BM expectations, with the 95% HPD interval for ω_x_ including zero, they revealed a pre-sympatric effect of biotic interactions on colonization. *Puya* lineages have smaller colonization rates in areas already occupied by phenotypically similar species, with probability greater than 95% (P(ω_1_ < 0) ≥ 0.95), consistent with competitive exclusion.

**Figure 4.**
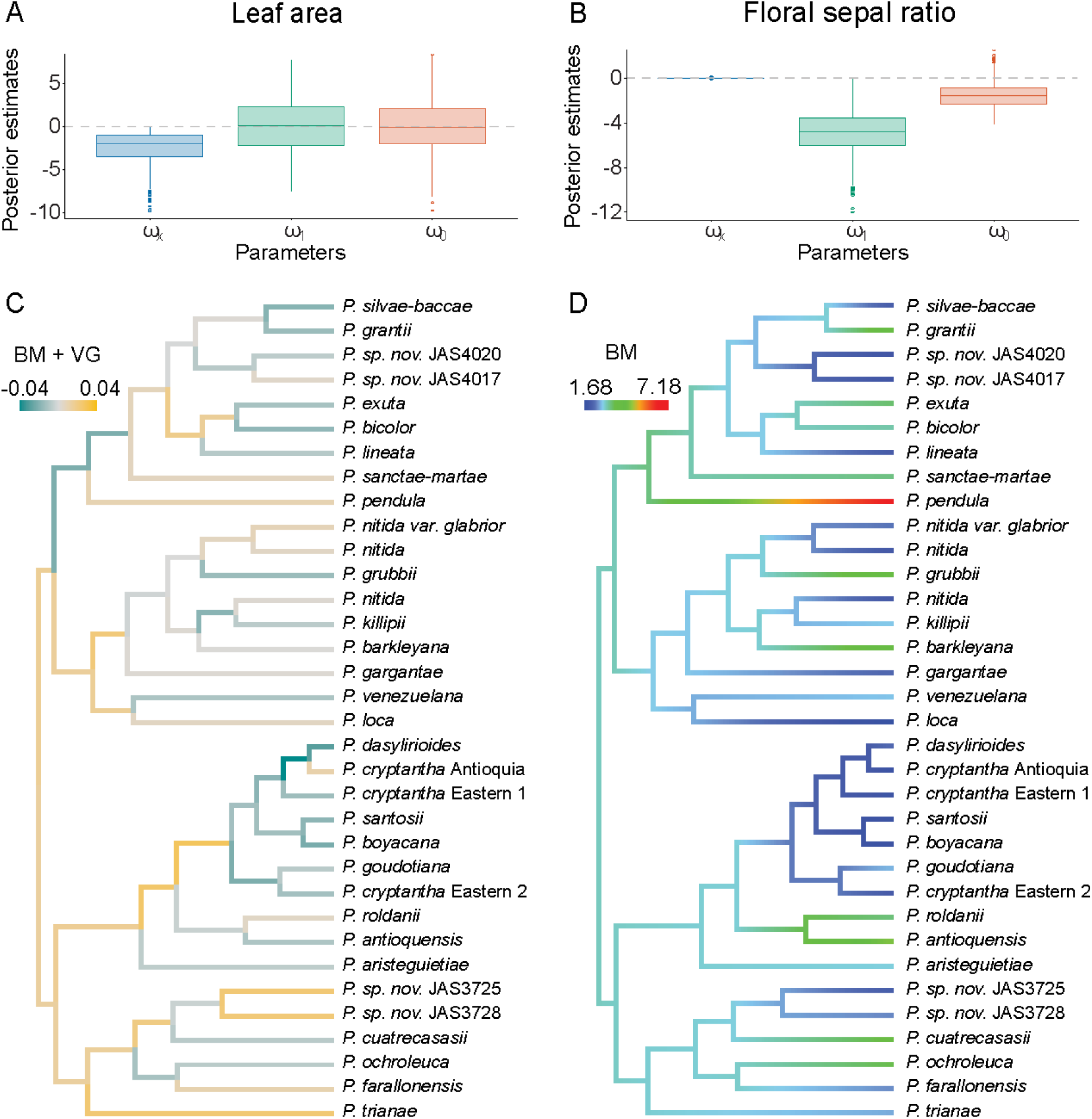
Effects of sympatry on trait evolution in the Septentrional Clade of *Puya*. Boxplots of posterior estimates for the parameters describing biotic interactions from the TRIBE model for each evaluated morphological trait. ωx indicates the effect on the character, ω1 the effect on colonization success, and ω0 on extirpation rates for (A) leaf area and (B) floral sepal ratio. (C) Leaf area evolution reconstruction under a branch-normalized signal-to-noise ratios of posterior jump distributions using a BM+VG model. Color scale indicates negative or positive deviations from the expectation of trait evolution explained predominantly under BM. (D) Floral sepal ratio evolution reconstruction using a BM model. Color scale indicates trait value.

### Trait evolution across the phylogenetic tree

We identified diverse phenotypic evolutionary patterns among the evaluated traits (Fig. 4C and D and SI Appendix, Fig. S10), showing multiple processes shaping the evolution of the phenotypes in this clade. For instance, floral bract ratio, floral sepal ratio, and plant length evolution were consistently described by a BM process (Fig. 4D and SI Appendix, Fig. S9 and Table S2). In contrast, leaf area and number of flowers were better explained by Lévy jump processes in most of the analyses that were performed (SI Appendix, Table S2), which indicates pulsed evolution.

For leaf area (Fig. 4C), we characterized the number of jumps under a model that describes pulsed evolution using a Variance Gamma process (VG) while including a BM component (BM+VG). We identified both positive and negative deviations from a BM expectation, explained by the model’s jump component. These deviations, or jumps, were identified across the recent radiation of *Puya*, including sympatric lineages. Regarding the number of flowers (SI Appendix, Fig. S10D), the model selected across independent runs varied; therefore, we present the estimated jump values under a model where pulsed evolution is described by a Jump Normal (JN) process (BM+JN). This is the simplest Lévy process implemented in the creepy-jerk software (Landis *et al*., 2013). We inferred strong jump values corresponding to positive deviations along the branches leading to *P. aristeguietae*, originally restricted to the Eastern Cordillera but later dispersing to the Cordillera de Mérida; *P. dasylirioides*, whose lineage recently dispersed to the Cordillera de Talamanca; and *P. goudotiana*, which evolved under sympatry in the Eastern Cordillera.

## Discussion

### Phenotypic divergence in sympatry: Evolution mode in *Puya* and inference of character displacement and competitive exclusion

The occurrence of morphological divergence in sympatry has been primarily studied in animals, while its prevalence in plants remains less understood. Recently, Olivares et al. (Olivares *et al*., 2025) conducted a comprehensive evaluation of conifers and palms and found no evidence of morphological divergence between sister taxa, in either sympatry or allopatry. The contrast between these predominantly widespread lowland lineages and *Puya* suggests that phenotypic divergence between sympatric species is more prevalent in geographically fragmented systems, such as sky-islands.

Our results show that the observed morphological divergence in *Puya* correlates with ecological and reproductive pressures associated with competitive exclusion and character displacement. Specifically, floral traits influence community assembly primarily through competitive exclusion, whereas vegetative traits appear more likely to diverge following sympatry through character displacement. Our finding that species with similar floral

morphologies are less likely to co-occur in the same location suggests that interspecific interactions centered on pollination specializations play a significant role in community assembly, even in the absence of post-sympatric divergence. Competitive pollination has been documented in other biodiversity hotspots, particularly in systems with limited pollinator availability, such as alpine, arid, and island environments (Sargent & Ackerly, 2008; Armbruster & Muchhala, 2009; Zhao *et al*., 2025). Although our dataset includes only a limited set of floral characters that are difficult to acquire through fieldwork and herbarium samples, these results point to an important, underappreciated role of floral competition in shaping community structure among *Puya* species.

Likewise, traits associated with character displacement are not only ecologically relevant but also closely tied to life history strategies in *Puya* and to its island-like ecological dynamics. For instance, the semelparous *P. goudotiana* shows gigantism, measured as large plant size and high flower production. The gigantism strategy can maximize reproductive output and increase seed dispersal success in extreme environmental constraints (Bodine *et al*., 2023; Aguirre-Santoro *et al*., 2024; Veldhuisen *et al*., 2024). Conversely, dwarf iteroparous species such as *P. brachystachya* and *P. silvae-baccae* specialize in ground-level pollinators and likely partition niches at a finer scale. Body size influences dispersal distances, effective population sizes, and the spatial grain at which organisms perceive their environment (Boucher *et al*., 2017), such that the evolution of both gigantism and dwarfism in island radiations represents alternative strategies for exploiting different ecological opportunities, with each size class potentially accessing distinct pollinator assemblages, microhabitats, or resource pools. Nürk et al. (Nürk *et al*., 2019) compared four oceanic and sky island radiations with their non-island relatives and found that plant height evolved into larger trait spaces in most island clades. These spaces were found to be rapidly saturated, a pattern that, as in *Puya*, may reflect an adaptive response.

We identified two distinct evolutionary pathways associated with character displacement: Brownian motion and pulsed evolution. On the one hand, plant length was best explained by a BM process (SI Appendix, Fig. S9), and the variance in this trait may have been sufficiently broad to allow phenotypic segregation. On the other hand, leaf area and flower number were better explained by a Lévy jump process, showing that evolutionary pulses explain their present-day variation. Simpson (Simpson, 1944) first proposed this process, pulsed evolution, as the result of lineage transitions between distinct adaptive zones that can lead to rapid periods of evolution, particularly when ecological and geographical opportunities are available. As *Puya* illustrates, such transitions may be driven by the same evolutionary pressures that promote phenotypic divergence via character displacement.

### The Septentrional Clade of *Puya* as a case of rapid diversification in Sky-island systems

Adaptive radiations demonstrate an intrinsic link between ecology and macroevolution and can therefore arise through multiple alternative pathways (Simpson, 1944; Givnish, 2015; Gillespie *et al*., 2020). However, three important aspects that we should consider when proposing a potential adaptive radiation are: the presence of ecological opportunity, elevated rates of lineage diversification, and adaptive phenotypic divergence (Gillespie *et al*., 2020). Here, we have shown that these three aspects are present in *Puya*’s Septentrional Clade.

In Bromeliads, the genus *Brocchinia* has been described as an adaptive radiation due to its diversity of nutrient-capture strategies, even though there is no evidence of accelerated diversification rates (Givnish *et al*., 2014; Givnish, 2015). Nonetheless, most plant lineages that diversified in the Andean páramo have shown fast diversification rates (Madriñán *et al*., 2013; Luebert & Weigend, 2014). In *Puya*, we identified an increase in net diversification rate that coincides with the geological period when the páramo ecosystem became available, suggesting that the availability of this ecosystem created ecological opportunities that promoted lineage diversification. This dynamic is shaped by both the topographic complexity of the Andes and the ecological constraints of the páramo ecosystem, where geographic and ecological isolation operate simultaneously to shape evolutionary trajectories (Flantua *et al*., 2019). The subsequent decrease in diversification rates that we identified may reflect a process of niche filling (Fig. 2) (Rabosky, 2009). Our biogeographic reconstruction shows that Colombia’s Eastern Cordillera served as the primary center of diversification for the Septentrional Clade. From there, *Puya* lineages expanded into other sky-islands primarily over the last two million years. This pattern aligns with those observed in other páramo radiations such as *Espeletia* (Pouchon *et al*., 2018, 2021), *Aragoa* (Bello *et al*., 2002), and *Linochilus* (Vargas *et al*., 2023), reinforcing the Eastern Cordillera’s role as both a cradle and corridor for páramo plant diversification.

Within páramo islands, we find a striking pattern of sympatric species divergence. Except for the Eastern Cordillera, most páramo islands host between one and five species that tend to either diverge morphologically when co-occurring or segregate elevationally when morphology is conserved (Fig. 1E). For instance, the Perijá Páramo island houses both *Puya silvae-baccae* (a dwarf species of ca. 10 cm) and *P. grantii* (an average-sized species of ca. 1 m), suggesting morphological divergence may be facilitating niche partitioning. Conversely, in Páramo de Belmira, *P. cryptantha* and *P. roldanii* share a similar morphology but are vertically stratified along elevation gradients. These findings support an “island rule” dynamic, where limited spatial and ecological resources promote either morphological divergence or fine-scale geographic partitioning (Buckley & Roughgarden, 2005). Moreover, empirical studies demonstrate that morphological variation accumulates more rapidly during the early phases of island colonization than during later stages, suggesting that the recent arrival of *Puya* lineages on these sky-islands may have created a temporal window for rapid phenotypic evolution (Nürk *et al*., 2019).

Overall, future pollination studies focusing on plant-animal interactions of *Puya* may reveal at a finer scale the mechanisms of rapid adaptation of this clade’s radiation. We propose that the recent diversification of the Septentrional Clade of *Puya* represents an example of how the Andes formation, together with interspecific interactions, has shaped the biodiversity observed in the tropics. Our work sheds new light on the relationship between sympatry and morphological divergence during plant macroevolution, demonstrating that continental sky-island systems can generate diversification patterns driven by ecological opportunity and interspecific interactions.

## Acknowledgments

We thank the curators and staff of the following herbaria for providing specimen access: B, COL, CUVC, F, FMB, HECASA, HUA, MA, MO, K, NY, P, and WU. N.C.R. and R.Z.F. were supported by the NSF with the award #DEB 2323170 Andean plant diversifications. J.A.S. and R.S.J. received funding from Colorado College for the generation of genomic data. Collection permits for all Colombian samples were granted by the Autoridad Nacional de Licencias Ambientales (ANLA) under permit number 2022095789-2-000.

## Competing interests

The authors declare no competing interests.

## Author contributions

N.C.R., J.A.S., and R.Z.F. designed research; N.C.R. and E.M.O. contributed new analytic tools; J.A.S. and R.S.J. contributed data; N.C.R. and E.M.O. analyzed data; N.C.R., J.A.S., and R.Z.F. wrote the paper; N.C.R., J.A.S., R.S.J., E.M.O., and R.Z.F. edited the manuscript.

## Data availability

Data and code are available on Zenodo DOI: 10.5281/zenodo.23963863.

## Supporting Information

**Fig. S1.** Species tree from the ASTRAL analysis.

**Fig. S2.** Time-calibrated Maximum Credibility Tree.

**Fig. S3**. Posterior estimates boxplots for cladogenetic parameters in the DEC+J model.

**Fig. S4**. Relationship between bract ratio and altitude.

**Fig. S5.** Relationship between sepal ratio and altitude.

**Fig. S6.** Relationship between leaf area and altitude.

**Fig. S7.** Relationship between flower number and altitude.

**Fig. S8.** Relationship between plant length and altitude.

**Fig. S9.** Boxplots of posterior estimates for the parameters describing biotic interactions from the TRIBE model for each evaluated morphological trait.

**Fig. S10.** Trait evolution in the Septentrional Clade of *Puya*.

**Table S1.** Posterior estimates for TRIBE analysis.

**Table S2**. Model comparison results of the pulsR analysis.

**Table S3.** Model comparison results of the BioGeoBEARS analysis.

**Dataset S1.** Morphological measurements and geographic occurrence data for species of the Septentrional Clade of *Puya*.

## Notes

### Competing Interest Statement

The authors have declared no competing interest.

## References

Aguirre-Santoro J, Zuluaga A, Stonesmyth E, Betancur J, Jabaily RS. 2024. Phylogenomics of Puya (Bromeliaceae): Evolution in the Andean slopes and sky island ecosystems. Journal of Systematics and Evolution 62: 257–274.

Armbruster WS, Muchhala N. 2009. Associations between floral specialization and species diversity: cause, effect, or correlation? Evolutionary Ecology 23: 159–179.

Bello MA, Chase MW, Olmstead RG, Rønsted N, Albach D. 2002. The Páramo Endemic Aragoa Is the Sister Genus of Plantago (Plantaginaceae; Lamiales): Evidence from Plastid rbcL and Nuclear Ribosomal ITS Sequence Data. Kew Bulletin 57: 585–597.

Bezanson J, Edelman A, Karpinski S, Shah VB. 2017. Julia: A fresh approach to numerical computing. SIAM Review 59: 65–98.

Bodine EN, Bush C, Capaldi A, Jabaily RS. 2023. Modelling differences in reproductive effort between iteroparous and semelparous reproductive strategies in Bromeliaceae. in silico Plants 5: diac019.

Boucher FC, Verboom GA, Musker S, Ellis AG. 2017. Plant size: a key determinant of diversification? The New Phytologist 216: 24–31.

Bouckaert R, Vaughan TG, Barido-Sottani J, Duchêne S, Fourment M, Gavryushkina A, Heled J, Jones G, Kühnert D, Maio ND, et al. 2019. BEAST 2.5: An advanced software platform for Bayesian evolutionary analysis. PLOS Computational Biology 15: e1006650.

Brown JH. 1971. Mammals on Mountaintops: Nonequilibrium Insular Biogeography. The American Naturalist 105: 467–478.

Brown J. 1978. The theory of insular biogeography and the distribution of boreal birds and mammals. Great Basin Naturalist Memoirs 2.

Brown WL, Wilson EO. 1956. Character displacement. Systematic Zoology 5: 49–64.

Buckley LB, Roughgarden J. 2005. Lizard habitat partitioning on islands: the interaction of local and landscape scales. Journal of Biogeography 32: 2113–2121.

Cortés AJ, Garzón LN, Valencia JB, Madriñán S. 2018. On the Causes of Rapid Diversification in the Páramos: Isolation by Ecology and Genomic Divergence in Espeletia. Frontiers in Plant Science 9: 1700.

Eisen KE, Geber MA. 2018. Ecological sorting and character displacement contribute to the structure of communities of Clarkia species. Journal of Evolutionary Biology 31: 1440–1458.

Flantua SGA, O’Dea A, Onstein RE, Giraldo C, Hooghiemstra H. 2019. The flickering connectivity system of the north Andean páramos. Journal of Biogeography 46: 1808–1825.

Gillespie RG, Bennett GM, De Meester L, Feder JL, Fleischer RC, Harmon LJ, Hendry AP, Knope ML, Mallet J, Martin C, et al. 2020. Comparing adaptive radiations across space, time, and taxa. The Journal of Heredity 111: 1–20.

Givnish TJ. 2015. Adaptive radiation versus ‘radiation’ and ‘explosive diversification’: why conceptual distinctions are fundamental to understanding evolution. New Phytologist 207: 297–303.

Givnish TJ, Barfuss MHJ, Van Ee B, Riina R, Schulte K, Horres R, Gonsiska PA, Jabaily RS, Crayn DM, Smith JAC, et al. 2011. Phylogeny, adaptive radiation, and historical biogeography in Bromeliaceae: insights from an eight-locus plastid phylogeny. American Journal of Botany 98: 872–895.

Givnish TJ, Barfuss MHJ, Van Ee B, Riina R, Schulte K, Horres R, Gonsiska PA, Jabaily RS, Crayn DM, Smith JAC, et al. 2014. Adaptive radiation, correlated and contingent evolution, and net species diversification in Bromeliaceae. Molecular Phylogenetics and Evolution 71: 55–78.

Guevara-Andino JE, Dávalos LM, Zapata F, Endara MJ, Cotoras DD, Chaves J, Claramunt S, López-Delgado J, Mendoza-Henao AM, Salazar-Valenzuela D, et al. 2025. Neotropics as a Cradle for Adaptive Radiations. Cold Spring Harbor Perspectives in Biology 17: a041452.

van der Hammen T. 1974. The Pleistocene changes of vegetation and climate in tropical South America. Journal of Biogeography 1: 3–26.

van der Hammen T, Cleef AM. 1986. Development of the high Andean páramo flora and vegetation. In: High altitude tropical biogeography. New York, NY: Oxford University Press, 153–201.

Höhna S. 2015. The time-dependent reconstructed evolutionary process with a key-role for mass-extinction events. Journal of Theoretical Biology 380: 321–331.

Höhna S, Landis MJ, Heath TA, Boussau B, Lartillot N, Moore BR, Huelsenbeck JP, Ronquist F. 2016. RevBayes: Bayesian phylogenetic inference using graphical models and an interactive model-specification language. Systematic Biology 65: 726–736.

Hooghiemstra H, Wijninga VM, Cleef AM. 2006. The paleobotanical record of Colombia: implications for biogeography and biodiversity. Annals of the Missouri Botanical Garden 93: 297–325.

Johnson MG, Pokorny L, Dodsworth S, Botigué LR, Cowan RS, Devault A, Eiserhardt WL, Epitawalage N, Forest F, Kim JT, et al. 2019. A Universal Probe Set for Targeted Sequencing of 353 Nuclear Genes from Any Flowering Plant Designed Using k-Medoids Clustering. Systematic Biology 68: 594–606.

Kalyaanamoorthy S, Minh BQ, Wong TKF, von Haeseler A, Jermiin LS. 2017. ModelFinder: fast model selection for accurate phylogenetic estimates. Nature Methods 14: 587–589.

Landis MJ, Freyman WA, Baldwin BG. 2018. Retracing the Hawaiian silversword radiation despite phylogenetic, biogeographic, and paleogeographic uncertainty. Evolution 72: 2343–2359.

Landis MJ, Schraiber JG. 2017. Pulsed evolution shaped modern vertebrate body sizes. Proceedings of the National Academy of Sciences of the United States of America 114: 13224–13229.

Landis MJ, Schraiber JG, Liang M. 2013. Phylogenetic analysis using Lévy processes: Finding jumps in the evolution of continuous traits. Systematic Biology 62: 193–204.

Londoño C, Cleef A, Madriñán S. 2014. Angiosperm flora and biogeography of the páramo region of Colombia, Northern Andes. Flora - Morphology, Distribution, Functional Ecology of Plants 209: 81–87.

Losos JB. 2010. Adaptive radiation, ecological opportunity, and evolutionary determinism. American Society of Naturalists E. O. Wilson award address. The American Naturalist 175: 623–639.

Losos JB, Glor RE, Kolbe JJ, Nicholson K. 2006. Adaptation, speciation, and convergence: A hierarchical analysis of adaptive radiation in Caribbean Anolis lizards. Annals of the Missouri Botanical Garden 93: 24–33.

Love SJ, Schweitzer JA, Woolbright SA, Bailey JK. 2023. Sky Islands Are a Global Tool for Predicting the Ecological and Evolutionary Consequences of Climate Change. Annual Review of Ecology, Evolution, and Systematics 54: 219–236.

Luebert F, Weigend M. 2014. Phylogenetic insights into Andean plant diversification. Frontiers in Ecology and Evolution 2.

Madriñán S, Cortés AJ, Richardson JE. 2013. Páramo is the world’s fastest evolving and coolest biodiversity hotspot. Frontiers in Genetics 4.

Magallon S, Sanderson MJ. 2001. Absolute diversification rates in angiosperm clades. Evolution 55: 1762–1780.

Magee AF, Höhna S, Vasylyeva TI, Leaché AD, Minin VN. 2020. Locally adaptive Bayesian birth-death model successfully detects slow and rapid rate shifts (DA Rasmussen, Ed.). PLOS Computational Biology 16: e1007999.

Maliet O, Hartig F, Morlon H. 2019. A model with many small shifts for estimating species-specific diversification rates. Nature Ecology & Evolution 3: 1086–1092.

Maliet O, Morlon H. 2022. Fast and Accurate Estimation of Species-Specific Diversification Rates Using Data Augmentation (J Rosindell, Ed.). Systematic Biology 71: 353–366.

Matzke NJ. 2013. Probabilistic historical biogeography: new models for founder-event speciation, imperfect detection, and fossils allow improved accuracy and model-testing. Frontiers of Biogeography 5.

Matzke NJ. 2014. Model Selection in Historical Biogeography Reveals that Founder-Event Speciation Is a Crucial Process in Island Clades. Systematic Biology 63: 951–970.

McGuire JA, Witt CC, Remsen JV, Corl A, Rabosky DL, Altshuler DL, Dudley R. 2014. Molecular Phylogenetics and the Diversification of Hummingbirds. Current Biology 24: 910–916.

Merrill RM, Dasmahapatra KK, Davey JW, Dell’Aglio DD, Hanly JJ, Huber B, Jiggins CD, Joron M, Kozak KM, Llaurens V, et al. 2015. The diversification of *Heliconius* butterflies: what have we learned in 150 years? Journal of Evolutionary Biology 28: 1417–1438.

Minh BQ, Schmidt HA, Chernomor O, Schrempf D, Woodhams MD, Von Haeseler A, Lanfear R. 2020. IQ-TREE 2: New Models and Efficient Methods for Phylogenetic Inference in the Genomic Era (E Teeling, Ed.). Molecular Biology and Evolution 37: 1530–1534.

Morita K, Yamamichi M. 2023. How does the magnitude of genetic variation affect ecological and reproductive character displacement? Population Ecology 65: 220–230.

Nürk NM, Atchison GW, Hughes CE. 2019. Island woodiness underpins accelerated disparification in plant radiations. The New Phytologist 224: 518–531.

Olivares I, Faurby S, Cámara-Leret R, Pigot AL. 2025. The likelihood of sympatric speciation and morphological divergence in plants. Proceedings of the National Academy of Sciences 122: e2508958122.

Ortiz EM, Höwener A, Shigita G, Raza M, Maurin O, Zuntini A, Forest F, Baker WJ, Schaefer H. 2026. A Novel Phylogenomics Pipeline Reveals Extensive Topological Conflict in the Evolution of the Angiosperm Order Cucurbitales (Y Yang, Ed.). Systematic Biology: syag046.

Pouchon C, Fernández A, Nassar JM, Boyer F, Aubert S, Lavergne S, Mavárez J. 2018. Phylogenomic Analysis of the Explosive Adaptive Radiation of the Espeletia Complex (Asteraceae) in the Tropical Andes. Systematic Biology 67: 1041–1060.

Pouchon C, Lavergne S, Fernández Á, Alberti A, Aubert S, Mavárez J. 2021. Phylogenetic signatures of ecological divergence and leapfrog adaptive radiation in Espeletia. American Journal of Botany 108: 113–128.

Quintero I, Landis MJ. 2020. Interdependent phenotypic and biogeographic evolution driven by biotic interactions. Systematic Biology 69: 739–755.

R Core Team. 2024. R: A Language and Environment for Statistical Computing. Vienna, Austria: R Foundation for Statistical Computing.

Rabosky DL. 2009. Ecological Limits on Clade Diversification in Higher Taxa. The American Naturalist 173: 662–674.

Revell LJ. 2012. phytools: an R package for phylogenetic comparative biology (and other things). Methods in Ecology and Evolution 3: 217–223.

Ronquist F, Sanmartín I. 2011. Phylogenetic Methods in Biogeography. Annual Review of Ecology, Evolution, and Systematics 42: 441–464.

Sanmartín I, Van Der Mark P, Ronquist F. 2008. Inferring dispersal: a Bayesian approach to phylogeny-based island biogeography, with special reference to the Canary Islands. Journal of Biogeography 35: 428–449.

Sargent RD, Ackerly DD. 2008. Plant–pollinator interactions and the assembly of plant communities. Trends in Ecology & Evolution 23: 123–130.

Seehausen O. 2006. African cichlid fish: a model system in adaptive radiation research. Proceedings of the Royal Society B: Biological Sciences 273: 1987–1998.

Simpson GG. 1944. Tempo and Mode in Evolution. Columbia University Press.

Sklenář P, Hedberg I, Cleef AM. 2014. Island biogeography of tropical alpine floras. Journal of Biogeography 41: 287–297.

Smith SA, Brown JW, Walker JF. 2018. So many genes, so little time: A practical approach to divergence-time estimation in the genomic era. PloS One 13: e0197433.

Stadler T. 2011. Mammalian phylogeny reveals recent diversification rate shifts. Proceedings of the National Academy of Sciences 108: 6187–6192.

Stroud JT, Delory BM, Barnes EM, Chase JM, De Meester L, Dieskau J, Grainger TN, Halliday FW, Kardol P, Knight TM, et al. 2024. Priority effects transcend scales and disciplines in biology. Trends in Ecology & Evolution 39: 677–688.

Stroud JT, Giery ST, Outerbridge M, Feeley KJ. 2019. Ecological character displacement alters the outcome of priority effects during community assembly. Ecology 100: e02727.

Vargas OM, Madriñán S, Simpson B. 2023. Allopatric speciation is more prevalent than parapatric ecological divergence in a recent high-Andean diversification (Linochilus: Asteraceae). PeerJ 11: e15479.

Veldhuisen L, Garcia Lino MC, Bodine E, Aguirre-Santoro J, Jabaily R. 2024. Description of Life History and Reproductive Size Thresholds in Three High Elevation Puya (Bromeliaceae). Selbyana 35: 1–10.

Vuilleumier F. 1970. Insular Biogeography in Continental Regions. I. The Northern Andes of South America. The American Naturalist 104: 373–388.

Witter MS, Carr GD. 1988. Adaptive radiation and genetic differentiation in the Hawaiian silversword alliance (Compositae: Madiinae). Evolution; International Journal of Organic Evolution 42: 1278–1287.

Zhang C, Scornavacca C, Molloy EK, Mirarab S. 2020. ASTRAL-Pro: Quartet-Based Species-Tree Inference despite Paralogy (J Thorne, Ed.). Molecular Biology and Evolution 37: 3292–3307.

Zhao C, Ba S, Zhang T, Li Y, Fang Q. 2025. Floral trait dissimilarity indirectly decreases heterospecific pollen transfer through pollinator sharing in a subalpine meadow. Oecologia 207: 55.

