## Supporting Information for "*Puya*: A new example of adaptive radiation in the Northern Andes"

### Figures

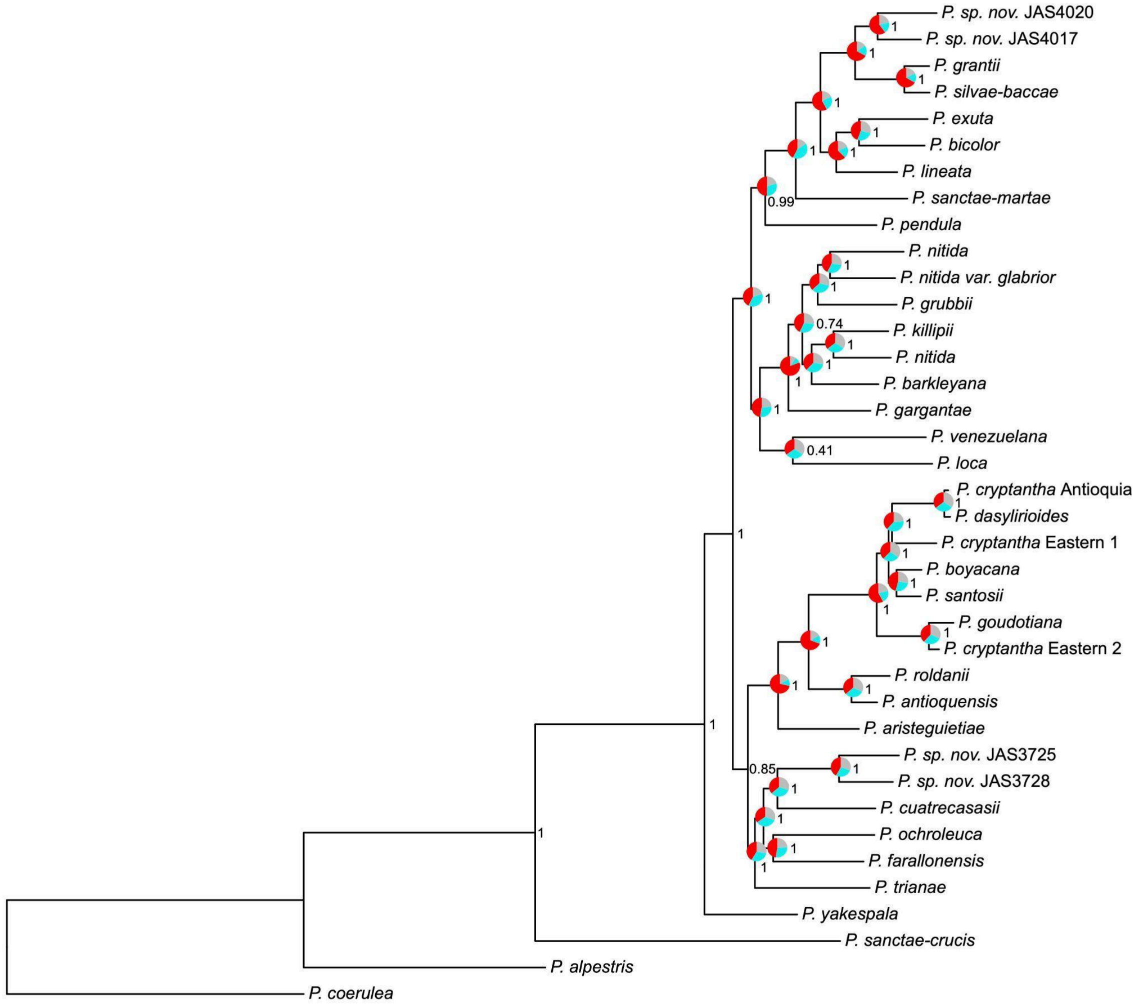


**Fig. S1.** Species tree from the ASTRAL analysis showing local posterior probabilities as numbers and quartet support as pie charts for each node. In the quartet support pies, red indicates the proportion supporting the dominant topology, blue indicates support for the best alternative topology, and grey represents support for the remaining topology.

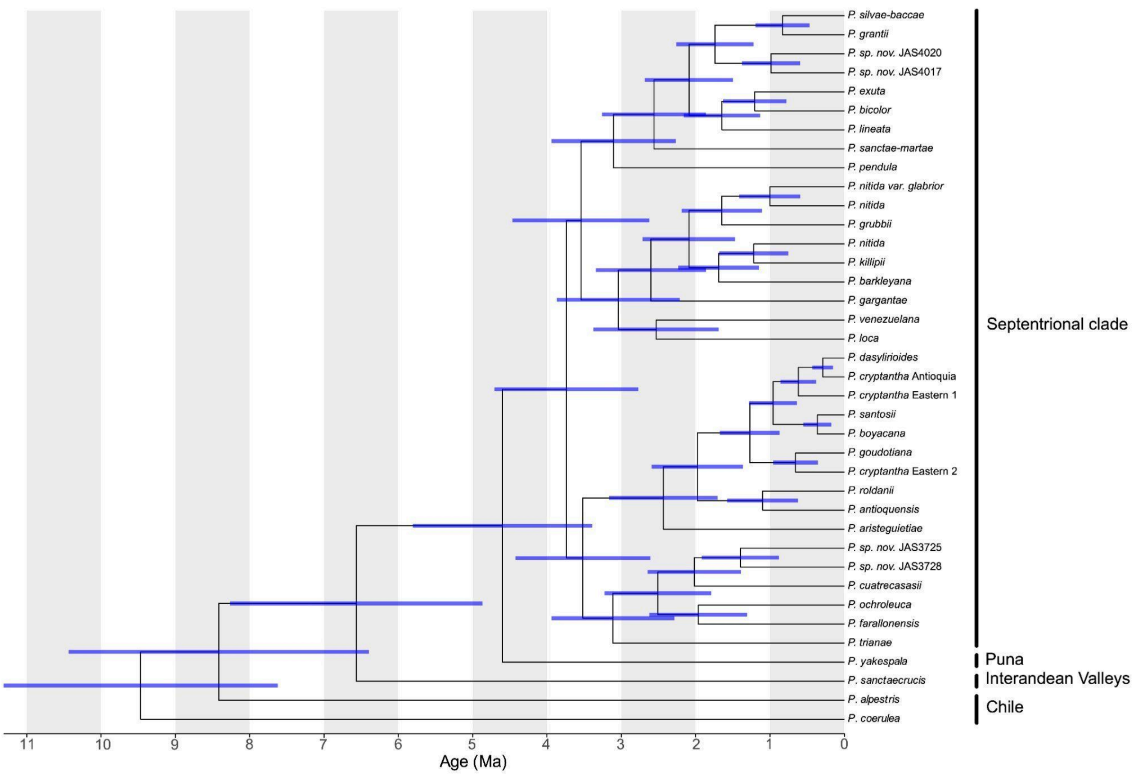


**Fig. S2.** Time-calibrated Maximum Credibility Tree (MCC) including the confidence intervals for the time estimations for each node.

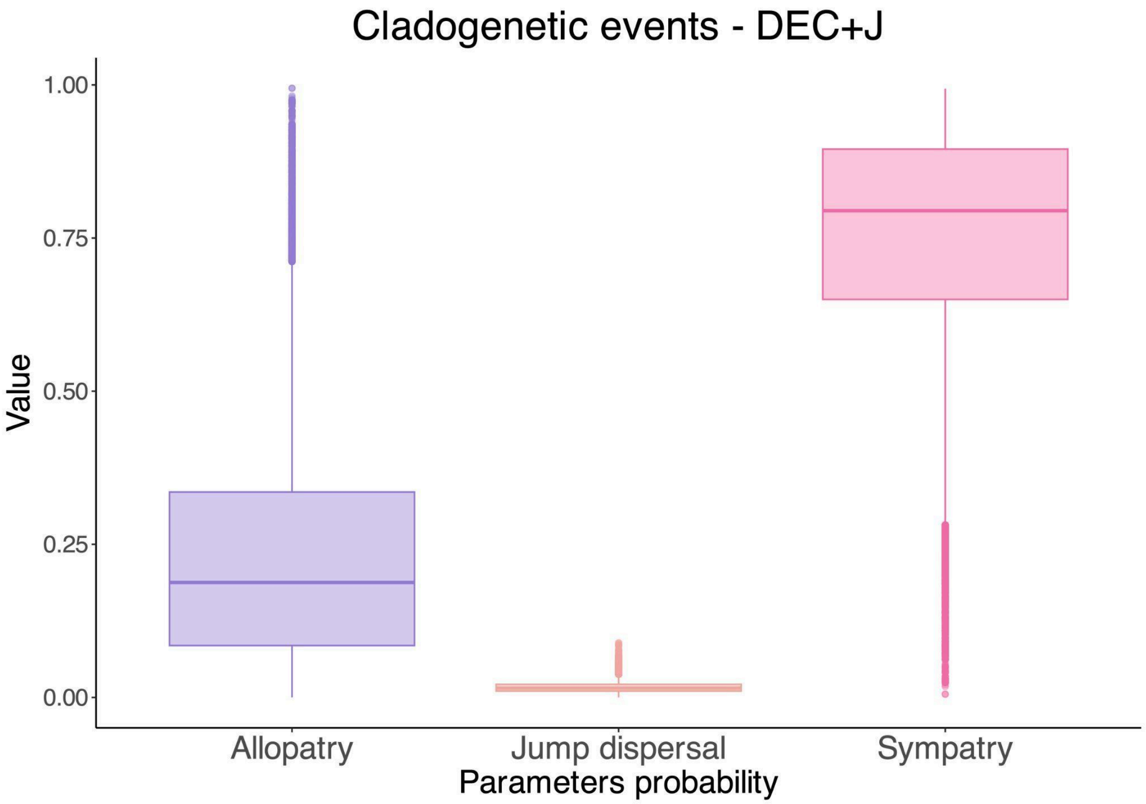


**Fig. S3.** Posterior estimates boxplots for cladogenetic parameters in the DEC+J model.

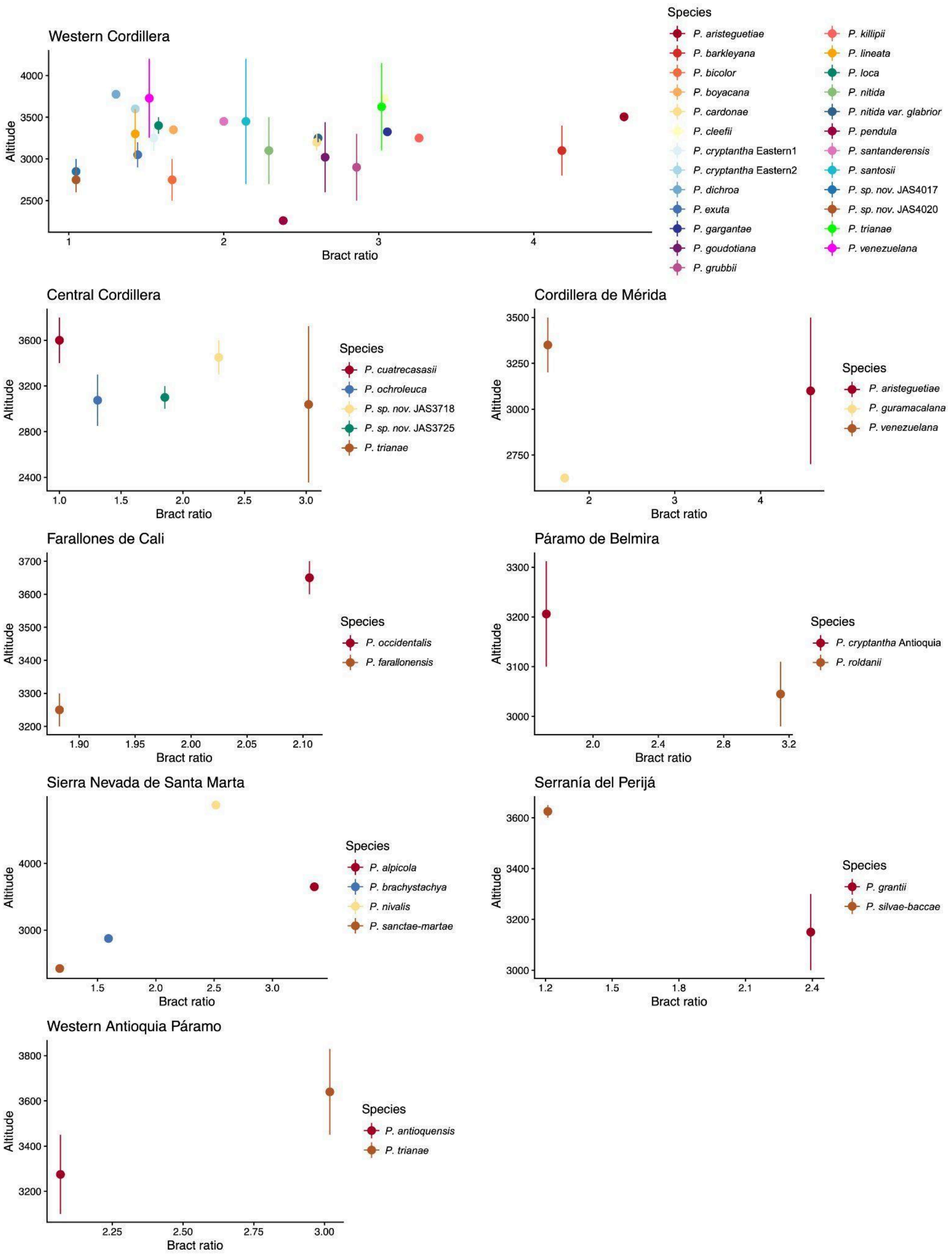


**Fig. S4.** Relationship between bract ratio and altitude assessed for *Puya* species inhabiting each of the considered areas. Dots represent mean phenotypic values, while lines indicate their respective ranges.

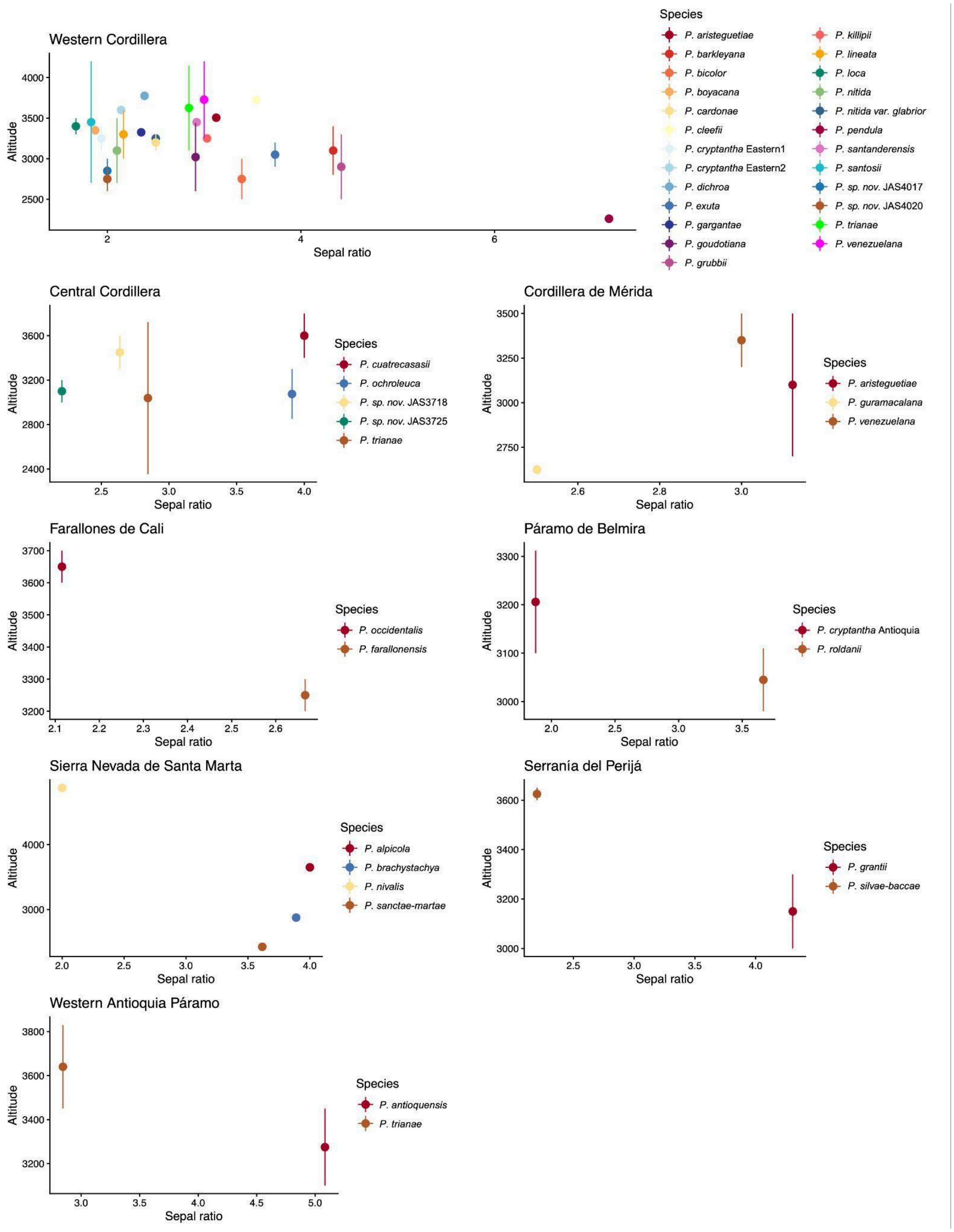


**Fig. S5.** Relationship between sepal ratio and altitude assessed for *Puya* species inhabiting each of the considered areas. Dots represent mean phenotypic values, while lines indicate their respective ranges.

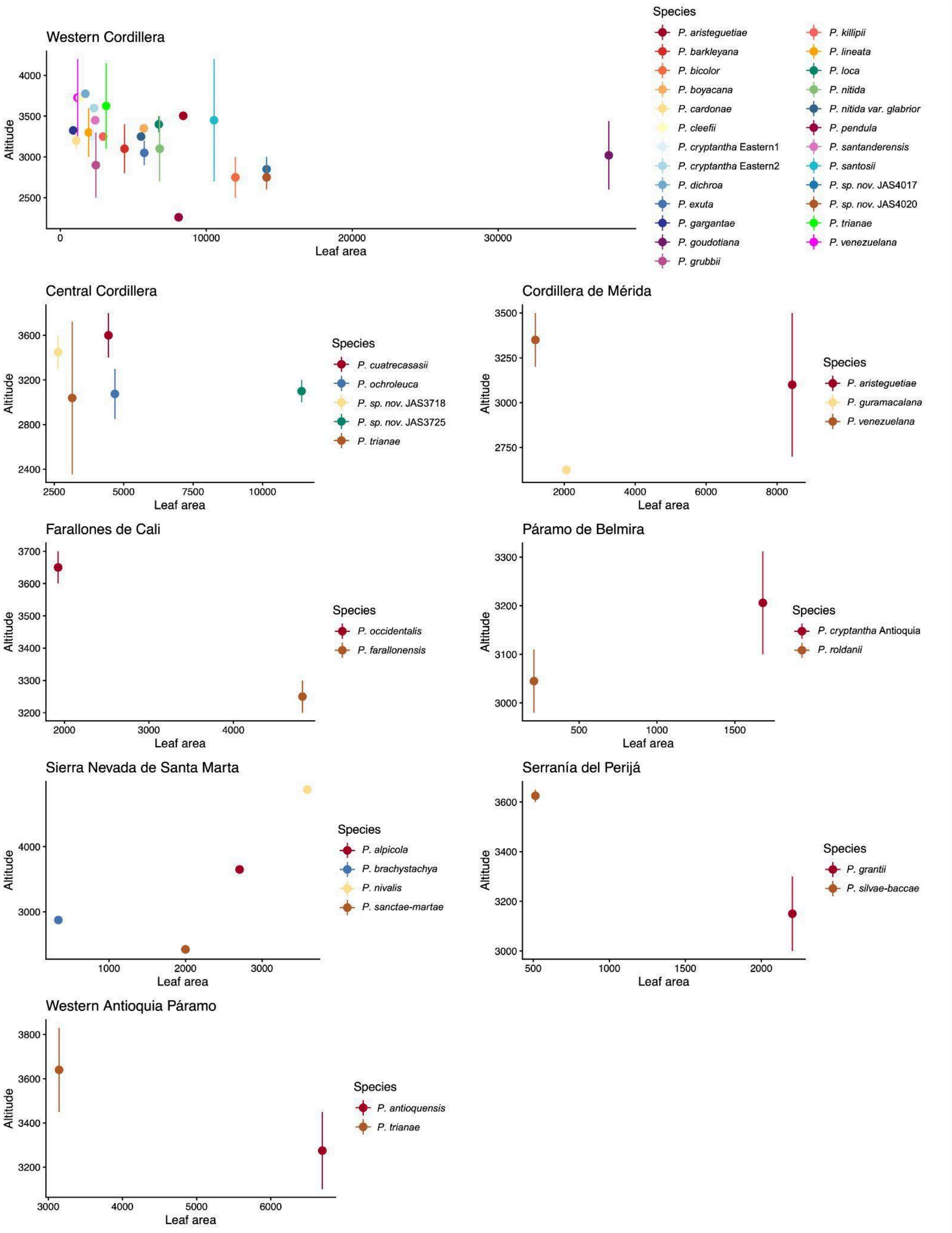


**Fig. S6.** Relationship between leaf area and altitude assessed for *Puya* species inhabiting each of the considered areas. Dots represent mean phenotypic values, while lines indicate their respective ranges.

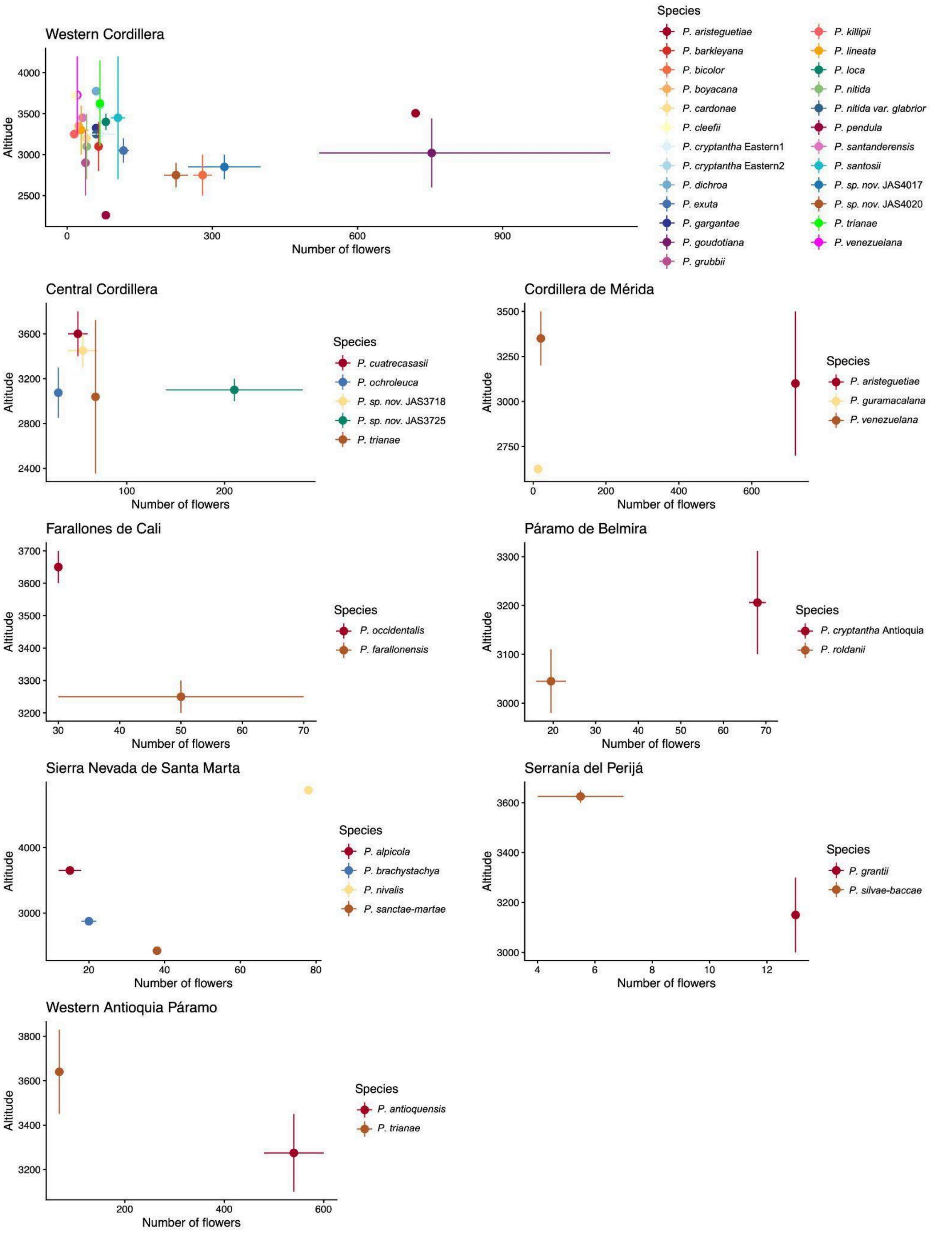


**Fig. S7.** Relationship between number of flowers and altitude assessed for *Puya* species inhabiting each of the considered areas. Dots represent mean phenotypic values, while lines indicate their respective ranges.

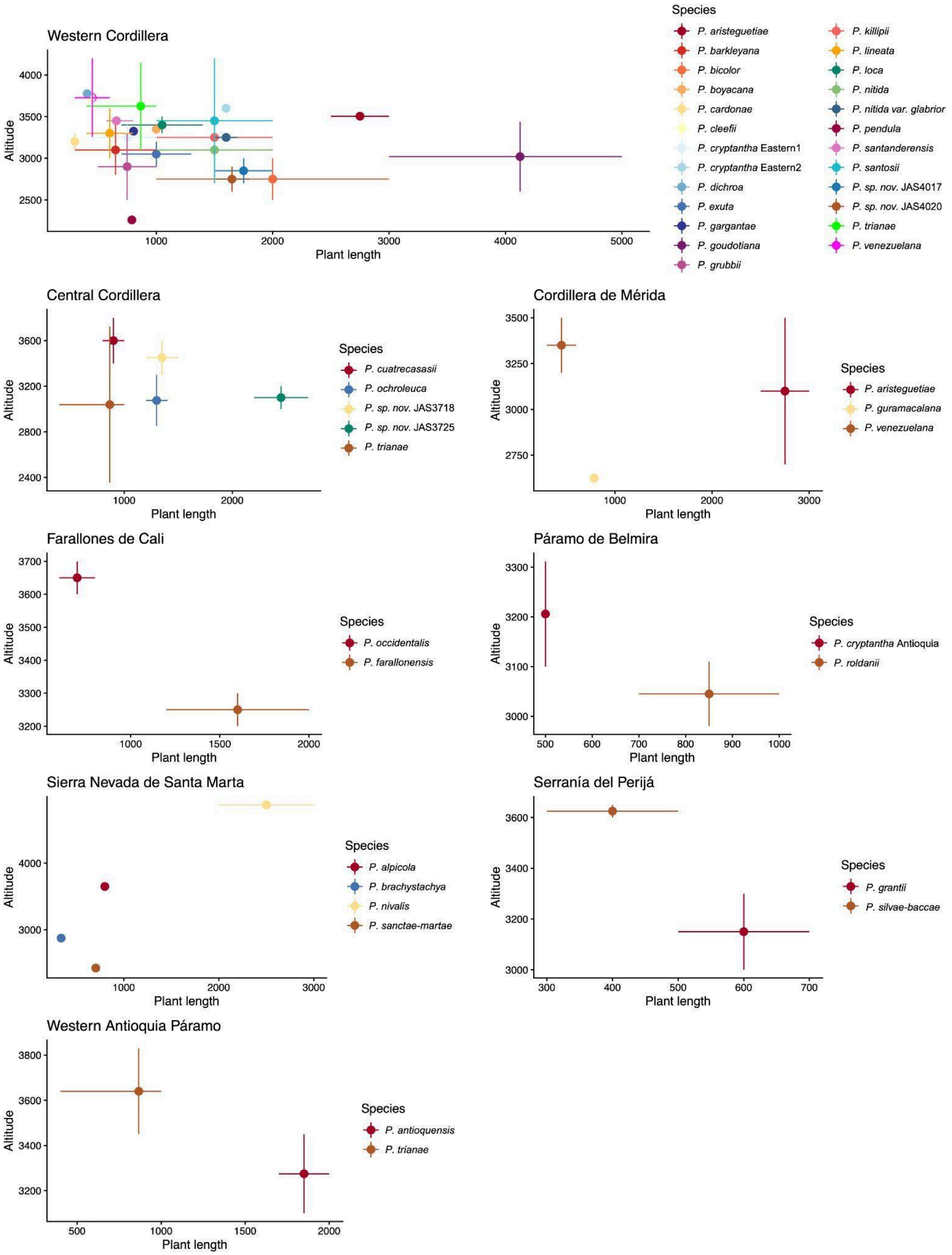


**Fig. S8.** Relationship between plant length and altitude assessed for *Puya* species inhabiting each of the considered areas. Dots represent mean phenotypic values, while lines indicate their respective ranges.

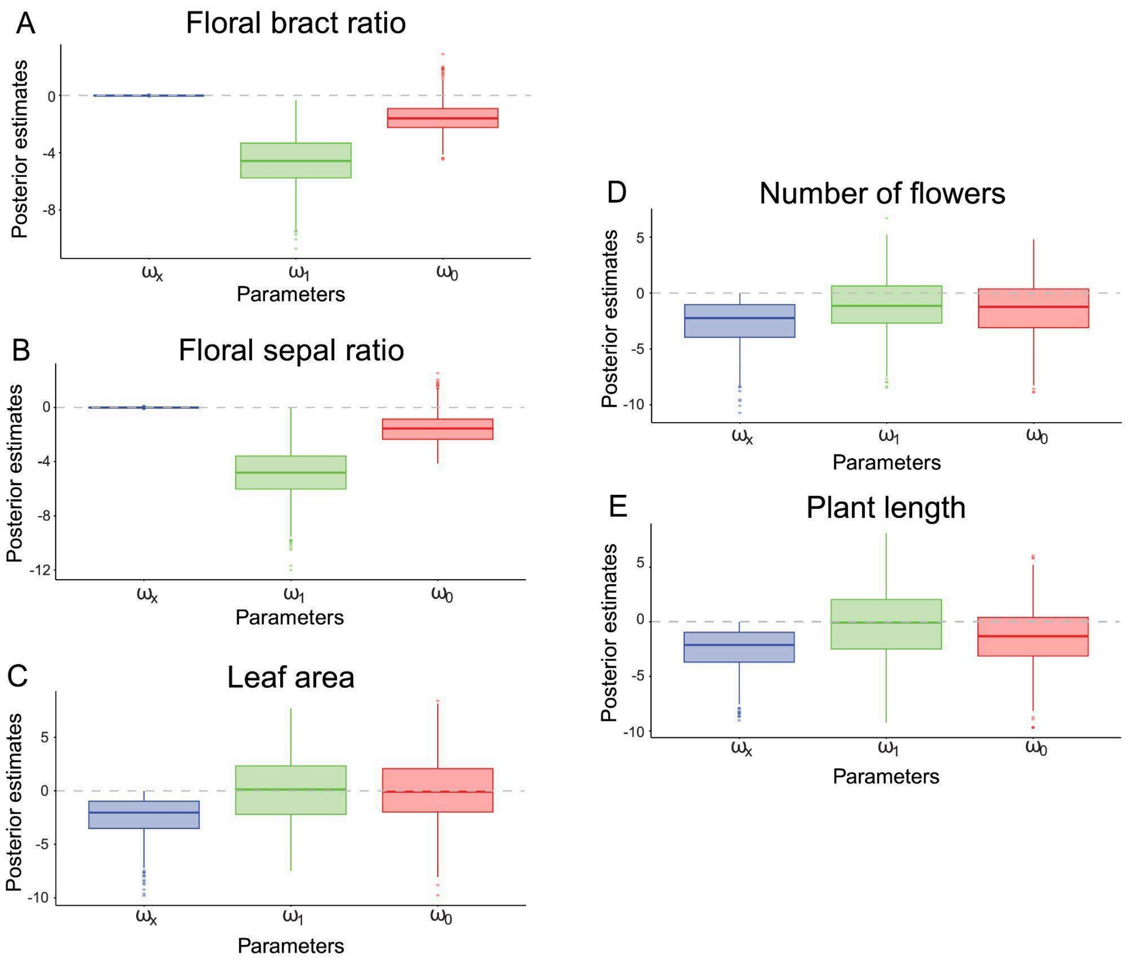


**Fig. S9.** Boxplots of posterior estimates for the parameters describing biotic interactions from the TRIBE model for each evaluated morphological trait. $\omega_x$ indicates the effect on the character, $\omega_1$ the effect on colonization success, and $\omega_0$ the effect on extirpation rates.

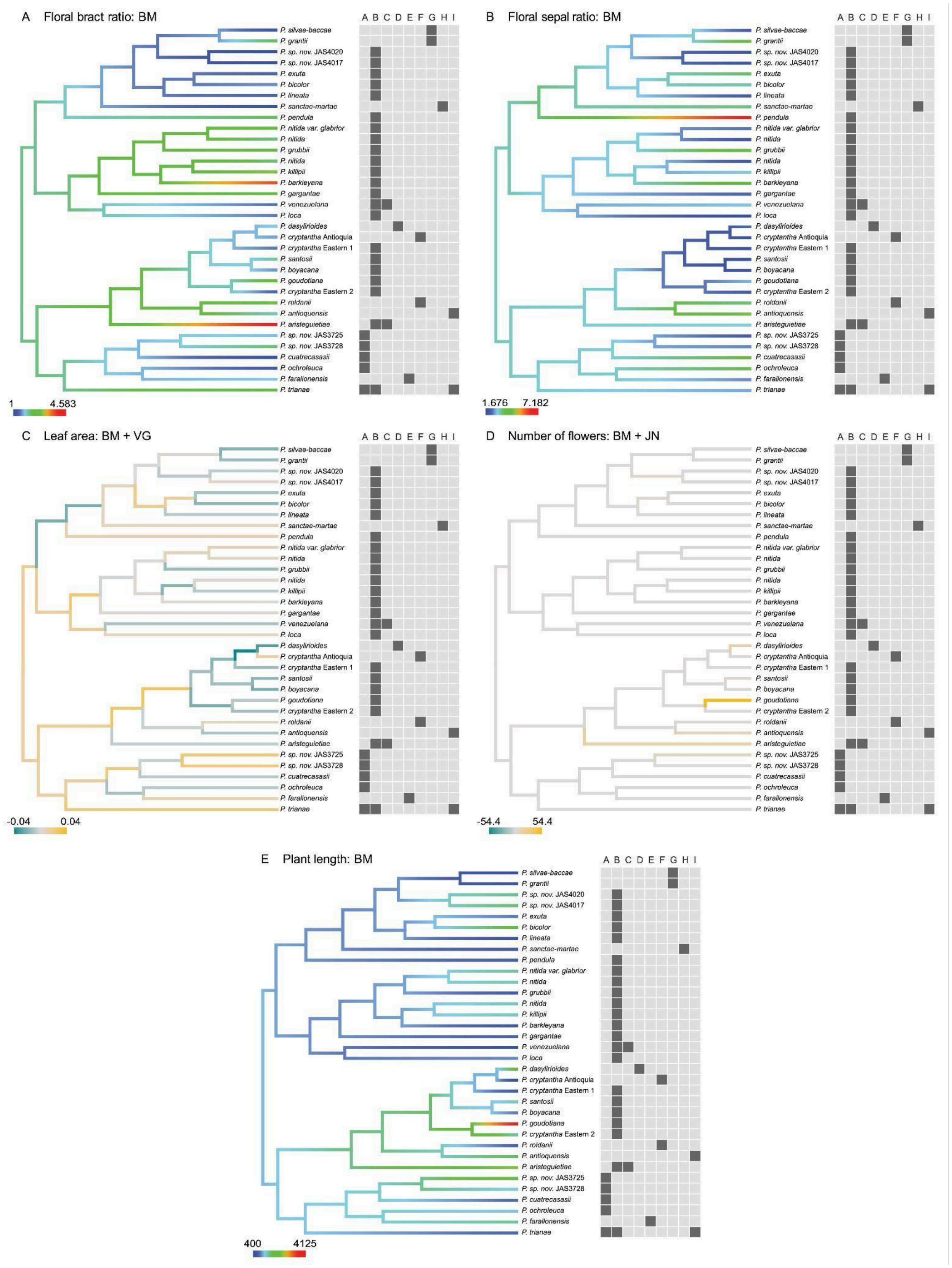


**Fig. S10.** Trait evolution in the Septentrional Clade of Puya. Floral bract ratio reconstructed under BM (A), floral sepal ratio reconstructed under BM (B), branch-normalized signal-to-noise ratios of posterior jump distributions for leaf area under BM+VG (C), branch-normalized signal-to-noise ratios of posterior jump distributions for number of flowers under BM+JN (D), and plant length reconstructed under BM (E). Colored panels adjacent to species names indicate species occurrence across the geographic regions examined in this study (area codes as in Fig. 1). In A, B, and E, colors represent trait values. In C and D, colors represent negative or positive deviations from trait evolution expected under a Brownian motion model.

**Tables**

**Table S1.** Posterior estimates for TRIBE analysis. Values correspond to their respective 95% highest probability density (HPD) interval.

| **Trait** | **Posterior median and 95% HPD estimates** $\omega_x$ | $\omega_1$ | $\omega_0$ |
|---|---|---|---|
| Floral bract ratio | -0.014 [-0.064, 0.023] | -4.615 [-8.345, -1.183] | -1.634 [-3.404, 0.808] |
| Floral sepal ratio | -0.005 [-0.051, 0.028] | -4.821 [-8.609, -1.122] | -1.589 [-3.699, 0.665] |
| Leaf area | -2.076 [-6.116, -0.011] | -0.098 [-5.344, 6.411] | 0.084 [-5.711, 5.288] |
| Number of flowers | -2.276 [-6.483, -0.004] | -1.147 [-5.961, 3.345] | -1.306 [-6.527, 3.141] |
| Plant length | -2.123 [-6.256, -0.002] | -0.122 [-7.164, 4.722] | -1.329 [-6.737, 3.626] |

**Table S2.** Model comparison results of the pulsR analysis. We conducted 10 independent parameter searches for each trait analyzed.

| Trait | Model | LogLikelihood | AIC | fOpt | k | AICc | Weight | Run |
|---|---|---|---|---|---|---|---|---|
| Plant length | BM | -192.33 | 388.66 | 192.33 | 2 | 389.05 | 0.45 | 1 |
| Plant length | EB | -191.56 | 389.12 | 191.56 | 3 | 389.92 | 0.29 | 1 |
| Plant length | JN | -192.33 | 390.66 | 192.33 | 3 | 391.46 | 0.13 | 1 |
| Plant length | OU | -192.34 | 390.67 | 192.34 | 3 | 391.47 | 0.13 | 1 |
| Plant length | BM | -192.33 | 388.66 | 192.33 | 2 | 389.05 | 0.53 | 2 |
| Plant length | EB | -192.33 | 390.66 | 192.33 | 3 | 391.46 | 0.16 | 2 |
| Plant length | OU | -192.33 | 390.66 | 192.33 | 3 | 391.46 | 0.16 | 2 |
| Plant length | NIG | -192.33 | 390.66 | 192.33 | 3 | 391.46 | 0.16 | 2 |
| Plant length | BM | -192.33 | 388.66 | 192.33 | 2 | 389.05 | 0.53 | 3 |
| Plant length | EB | -192.33 | 390.66 | 192.33 | 3 | 391.46 | 0.16 | 3 |
| Plant length | VG | -192.33 | 390.66 | 192.33 | 3 | 391.46 | 0.16 | 3 |
| Plant length | OU | -192.37 | 390.73 | 192.37 | 3 | 391.53 | 0.15 | 3 |

| Trait | Model | LogLikelihood | AIC | fOpt | k | AICc | Weight | Run |
|---|---|---|---|---|---|---|---|---|
| Plant length | BM | -192.33 | 388.66 | 192.33 | 2 | 389.05 | 0.53 | 4 |
| Plant length | EB | -192.33 | 390.66 | 192.33 | 3 | 391.46 | 0.16 | 4 |
| Plant length | NIG | -192.33 | 390.66 | 192.33 | 3 | 391.46 | 0.16 | 4 |
| Plant length | OU | -192.38 | 390.75 | 192.38 | 3 | 391.55 | 0.15 | 4 |
| Plant length | BM | -192.33 | 388.66 | 192.33 | 2 | 389.05 | 0.45 | 5 |
| Plant length | EB | -191.56 | 389.12 | 191.56 | 3 | 389.92 | 0.29 | 5 |
| Plant length | VG | -192.33 | 390.66 | 192.33 | 3 | 391.46 | 0.13 | 5 |
| Plant length | OU | -192.33 | 390.66 | 192.33 | 3 | 391.46 | 0.13 | 5 |
| Plant length | BM | -192.33 | 388.66 | 192.33 | 2 | 389.05 | 0.45 | 6 |
| Plant length | EB | -191.56 | 389.12 | 191.56 | 3 | 389.92 | 0.29 | 6 |
| Plant length | OU | -192.33 | 390.66 | 192.33 | 3 | 391.46 | 0.13 | 6 |
| Plant length | NIG | -192.33 | 390.66 | 192.33 | 3 | 391.46 | 0.13 | 6 |
| Plant length | BM | -192.33 | 388.66 | 192.33 | 2 | 389.05 | 0.45 | 7 |
| Plant length | EB | -191.56 | 389.12 | 191.56 | 3 | 389.92 | 0.29 | 7 |

| Trait | Model | LogLikelihood | AIC | fOpt | k | AICc | Weight | Run |
|---|---|---|---|---|---|---|---|---|
| Plant length | NIG | -192.33 | 390.67 | 192.33 | 3 | 391.47 | 0.13 | 7 |
| Plant length | OU | -192.38 | 390.75 | 192.38 | 3 | 391.55 | 0.13 | 7 |
| Plant length | BM | -192.33 | 388.66 | 192.33 | 2 | 389.05 | 0.45 | 8 |
| Plant length | EB | -191.56 | 389.12 | 191.56 | 3 | 389.92 | 0.29 | 8 |
| Plant length | VG | -192.33 | 390.66 | 192.33 | 3 | 391.46 | 0.13 | 8 |
| Plant length | OU | -192.38 | 390.75 | 192.38 | 3 | 391.55 | 0.13 | 8 |
| Plant length | BM | -192.33 | 388.66 | 192.33 | 2 | 389.05 | 0.45 | 9 |
| Plant length | EB | -191.56 | 389.12 | 191.56 | 3 | 389.92 | 0.29 | 9 |
| Plant length | NIG | -192.33 | 390.66 | 192.33 | 3 | 391.46 | 0.13 | 9 |
| Plant length | OU | -192.38 | 390.75 | 192.38 | 3 | 391.55 | 0.13 | 9 |
| Plant length | BM | -192.33 | 388.66 | 192.33 | 2 | 389.05 | 0.45 | 10 |
| Plant length | EB | -191.56 | 389.12 | 191.56 | 3 | 389.92 | 0.29 | 10 |
| Plant length | VG | -192.33 | 390.66 | 192.33 | 3 | 391.46 | 0.13 | 10 |
| Plant length | OU | -192.33 | 390.66 | 192.33 | 3 | 391.46 | 0.13 | 10 |

| Trait | Model | LogLikelihood | AIC | fOpt | k | AICc | Weight | Run |
|---|---|---|---|---|---|---|---|---|
| Leaf area | BMVG | -184.30 | 376.59 | 184.30 | 4 | 377.97 | 0.70 | 1 |
| Leaf area | BM | -188.13 | 380.26 | 188.13 | 2 | 380.65 | 0.18 | 1 |
| Leaf area | OU | -188.13 | 382.26 | 188.13 | 3 | 383.06 | 0.06 | 1 |
| Leaf area | EB | -188.13 | 382.26 | 188.13 | 3 | 383.06 | 0.06 | 1 |
| Leaf area | BMVG | -184.30 | 376.59 | 184.30 | 4 | 377.97 | 0.70 | 2 |
| Leaf area | BM | -188.13 | 380.26 | 188.13 | 2 | 380.65 | 0.18 | 2 |
| Leaf area | EB | -188.13 | 382.26 | 188.13 | 3 | 383.06 | 0.06 | 2 |
| Leaf area | OU | -188.13 | 382.26 | 188.13 | 3 | 383.06 | 0.06 | 2 |
| Leaf area | BMVG | -184.30 | 376.59 | 184.30 | 4 | 377.97 | 0.70 | 3 |
| Leaf area | BM | -188.13 | 380.26 | 188.13 | 2 | 380.65 | 0.18 | 3 |
| Leaf area | OU | -188.13 | 382.26 | 188.13 | 3 | 383.06 | 0.06 | 3 |
| Leaf area | EB | -188.13 | 382.26 | 188.13 | 3 | 383.06 | 0.06 | 3 |
| Leaf area | BM | -188.13 | 380.26 | 188.13 | 2 | 380.65 | 0.53 | 4 |
| Leaf area | VG | -188.13 | 382.26 | 188.13 | 3 | 383.06 | 0.16 | 4 |

| Trait | Model | LogLikelihood | AIC | fOpt | k | AICc | Weight | Run |
|---|---|---|---|---|---|---|---|---|
| Leaf area | OU | -188.13 | 382.26 | 188.13 | 3 | 383.06 | 0.16 | 4 |
| Leaf area | EB | -188.13 | 382.26 | 188.13 | 3 | 383.06 | 0.16 | 4 |
| Leaf area | BMVG | -184.30 | 376.59 | 184.30 | 4 | 377.97 | 0.70 | 5 |
| Leaf area | BM | -188.13 | 380.26 | 188.13 | 2 | 380.65 | 0.18 | 5 |
| Leaf area | EB | -188.13 | 382.26 | 188.13 | 3 | 383.06 | 0.06 | 5 |
| Leaf area | OU | -188.13 | 382.26 | 188.13 | 3 | 383.06 | 0.06 | 5 |
| Leaf area | BM | -188.13 | 380.26 | 188.13 | 2 | 380.65 | 0.53 | 6 |
| Leaf area | VG | -188.13 | 382.26 | 188.13 | 3 | 383.06 | 0.16 | 6 |
| Leaf area | EB | -188.13 | 382.26 | 188.13 | 3 | 383.06 | 0.16 | 6 |
| Leaf area | OU | -188.13 | 382.26 | 188.13 | 3 | 383.06 | 0.16 | 6 |
| Leaf area | BM | -188.13 | 380.26 | 188.13 | 2 | 380.65 | 0.53 | 7 |
| Leaf area | EB | -188.13 | 382.26 | 188.13 | 3 | 383.06 | 0.16 | 7 |
| Leaf area | OU | -188.13 | 382.26 | 188.13 | 3 | 383.06 | 0.16 | 7 |
| Leaf area | JN | -188.13 | 382.26 | 188.13 | 3 | 383.06 | 0.16 | 7 |

| Trait | Model | LogLikelihood | AIC | fOpt | k | AICc | Weight | Run |
|---|---|---|---|---|---|---|---|---|
| Leaf area | BMVG | -184.30 | 376.59 | 184.30 | 4 | 377.97 | 0.70 | 8 |
| Leaf area | BM | -188.13 | 380.26 | 188.13 | 2 | 380.65 | 0.18 | 8 |
| Leaf area | EB | -188.13 | 382.26 | 188.13 | 3 | 383.06 | 0.06 | 8 |
| Leaf area | OU | -188.13 | 382.26 | 188.13 | 3 | 383.06 | 0.06 | 8 |
| Leaf area | VG | -184.30 | 374.59 | 184.30 | 3 | 375.39 | 0.90 | 9 |
| Leaf area | BM | -188.13 | 380.26 | 188.13 | 2 | 380.65 | 0.06 | 9 |
| Leaf area | EB | -188.13 | 382.26 | 188.13 | 3 | 383.06 | 0.02 | 9 |
| Leaf area | OU | -188.13 | 382.26 | 188.13 | 3 | 383.06 | 0.02 | 9 |
| Leaf area | BMVG | -184.30 | 376.59 | 184.30 | 4 | 377.97 | 0.70 | 10 |
| Leaf area | BM | -188.13 | 380.26 | 188.13 | 2 | 380.65 | 0.18 | 10 |
| Leaf area | OU | -188.13 | 382.26 | 188.13 | 3 | 383.06 | 0.06 | 10 |
| Leaf area | EB | -188.13 | 382.26 | 188.13 | 3 | 383.06 | 0.06 | 10 |
| Flower number | BMVG | -138.47 | 284.94 | 138.47 | 4 | 286.32 | 0.98 | 1 |
| Flower number | BM | -145.16 | 294.33 | 145.16 | 2 | 294.72 | 0.01 | 1 |

| Trait | Model | LogLikelihood | AIC | fOpt | k | AICc | Weight | Run |
|---|---|---|---|---|---|---|---|---|
| Flower number | EB | -145.16 | 296.33 | 145.16 | 3 | 297.13 | 0.00 | 1 |
| Flower number | OU | -145.16 | 296.33 | 145.16 | 3 | 297.13 | 0.00 | 1 |
| Flower number | BM | -145.16 | 294.33 | 145.16 | 2 | 294.72 | 0.57 | 2 |
| Flower number | EB | -144.79 | 295.58 | 144.79 | 3 | 296.38 | 0.25 | 2 |
| Flower number | OU | -145.39 | 296.78 | 145.39 | 3 | 297.58 | 0.14 | 2 |
| Flower number | BMJN | -145.16 | 298.33 | 145.16 | 4 | 299.71 | 0.05 | 2 |
| Flower number | NIG | -139.85 | 285.70 | 139.85 | 3 | 286.50 | 0.98 | 3 |
| Flower number | BM | -145.16 | 294.33 | 145.16 | 2 | 294.72 | 0.02 | 3 |
| Flower number | EB | -145.16 | 296.33 | 145.16 | 3 | 297.13 | 0.00 | 3 |
| Flower number | OU | -145.39 | 296.78 | 145.39 | 3 | 297.58 | 0.00 | 3 |
| Flower number | BMVG | -138.47 | 284.94 | 138.47 | 4 | 286.32 | 0.98 | 4 |
| Flower number | BM | -145.16 | 294.33 | 145.16 | 2 | 294.72 | 0.01 | 4 |
| Flower number | EB | -145.16 | 296.33 | 145.16 | 3 | 297.13 | 0.00 | 4 |
| Flower number | OU | -145.39 | 296.78 | 145.39 | 3 | 297.58 | 0.00 | 4 |

| Trait | Model | LogLikelihood | AIC | fOpt | k | AICc | Weight | Run |
|---|---|---|---|---|---|---|---|---|
| Flower number | BM | -145.16 | 294.33 | 145.16 | 2 | 294.72 | 0.49 | 5 |
| Flower number | EB | -144.79 | 295.58 | 144.79 | 3 | 296.38 | 0.21 | 5 |
| Flower number | NIG | -145.16 | 296.33 | 145.16 | 3 | 297.13 | 0.15 | 5 |
| Flower number | OU | -145.17 | 296.33 | 145.17 | 3 | 297.13 | 0.15 | 5 |
| Flower number | BMNIG | -139.85 | 287.70 | 139.85 | 4 | 289.08 | 0.91 | 6 |
| Flower number | BM | -145.16 | 294.33 | 145.16 | 2 | 294.72 | 0.05 | 6 |
| Flower number | EB | -145.16 | 296.33 | 145.16 | 3 | 297.13 | 0.02 | 6 |
| Flower number | OU | -145.16 | 296.33 | 145.16 | 3 | 297.13 | 0.02 | 6 |
| Flower number | BM | -145.16 | 294.33 | 145.16 | 2 | 294.72 | 0.49 | 7 |
| Flower number | EB | -144.79 | 295.58 | 144.79 | 3 | 296.38 | 0.21 | 7 |
| Flower number | NIG | -145.16 | 296.33 | 145.16 | 3 | 297.13 | 0.15 | 7 |
| Flower number | OU | -145.16 | 296.33 | 145.16 | 3 | 297.13 | 0.15 | 7 |
| Flower number | NIG | -139.85 | 285.70 | 139.85 | 3 | 286.50 | 0.98 | 8 |
| Flower number | BM | -145.16 | 294.33 | 145.16 | 2 | 294.72 | 0.02 | 8 |

| Trait | Model | LogLikelihood | AIC | fOpt | k | AICc | Weight | Run |
|---|---|---|---|---|---|---|---|---|
| Flower number | EB | -145.16 | 296.33 | 145.16 | 3 | 297.13 | 0.00 | 8 |
| Flower number | OU | -145.39 | 296.78 | 145.39 | 3 | 297.58 | 0.00 | 8 |
| Flower number | BM | -145.16 | 294.33 | 145.16 | 2 | 294.72 | 0.55 | 9 |
| Flower number | EB | -144.79 | 295.58 | 144.79 | 3 | 296.38 | 0.24 | 9 |
| Flower number | OU | -145.16 | 296.33 | 145.16 | 3 | 297.13 | 0.16 | 9 |
| Flower number | BMNIG | -145.16 | 298.33 | 145.16 | 4 | 299.71 | 0.05 | 9 |
| Flower number | NIG | -139.85 | 285.70 | 139.85 | 3 | 286.50 | 0.97 | 10 |
| Flower number | BM | -145.16 | 294.33 | 145.16 | 2 | 294.72 | 0.02 | 10 |
| Flower number | EB | -144.79 | 295.58 | 144.79 | 3 | 296.38 | 0.01 | 10 |
| Flower number | OU | -145.38 | 296.75 | 145.38 | 3 | 297.55 | 0.00 | 10 |
| Bract ratio | BM | -39.96 | 83.92 | 39.96 | 2 | 84.31 | 0.52 | 1 |
| Bract ratio | EB | -39.91 | 85.83 | 39.91 | 3 | 86.63 | 0.16 | 1 |
| Bract ratio | OU | -39.96 | 85.92 | 39.96 | 3 | 86.72 | 0.16 | 1 |
| Bract ratio | JN | -39.97 | 85.94 | 39.97 | 3 | 86.74 | 0.16 | 1 |
| Bract ratio | BM | -39.96 | 83.92 | 39.96 | 2 | 84.31 | 0.52 | 2 |

| Trait | Model | LogLikelihood | AIC | fOpt | k | AICc | Weight | Run |
|---|---|---|---|---|---|---|---|---|
| Bract ratio | EB | -39.91 | 85.83 | 39.91 | 3 | 86.63 | 0.16 | 2 |
| Bract ratio | VG | -39.96 | 85.92 | 39.96 | 3 | 86.72 | 0.16 | 2 |
| Bract ratio | OU | -39.96 | 85.92 | 39.96 | 3 | 86.72 | 0.16 | 2 |
| Bract ratio | BM | -39.96 | 83.92 | 39.96 | 2 | 84.31 | 0.52 | 3 |
| Bract ratio | EB | -39.91 | 85.83 | 39.91 | 3 | 86.63 | 0.16 | 3 |
| Bract ratio | NIG | -39.96 | 85.92 | 39.96 | 3 | 86.72 | 0.16 | 3 |
| Bract ratio | OU | -39.96 | 85.92 | 39.96 | 3 | 86.72 | 0.16 | 3 |
| Bract ratio | BM | -39.96 | 83.92 | 39.96 | 2 | 84.31 | 0.53 | 4 |
| Bract ratio | EB | -39.96 | 85.92 | 39.96 | 3 | 86.72 | 0.16 | 4 |
| Bract ratio | NIG | -39.96 | 85.92 | 39.96 | 3 | 86.72 | 0.16 | 4 |
| Bract ratio | OU | -39.96 | 85.92 | 39.96 | 3 | 86.72 | 0.16 | 4 |
| Bract ratio | BM | -39.96 | 83.92 | 39.96 | 2 | 84.31 | 0.53 | 5 |
| Bract ratio | EB | -39.96 | 85.92 | 39.96 | 3 | 86.72 | 0.16 | 5 |
| Bract ratio | OU | -39.96 | 85.92 | 39.96 | 3 | 86.72 | 0.16 | 5 |
| Bract ratio | NIG | -39.96 | 85.92 | 39.96 | 3 | 86.72 | 0.16 | 5 |
| Bract ratio | BM | -39.96 | 83.92 | 39.96 | 2 | 84.31 | 0.53 | 6 |
| Bract ratio | EB | -39.96 | 85.92 | 39.96 | 3 | 86.72 | 0.16 | 6 |
| Bract ratio | OU | -39.96 | 85.92 | 39.96 | 3 | 86.72 | 0.16 | 6 |
| Bract ratio | VG | -39.96 | 85.92 | 39.96 | 3 | 86.72 | 0.16 | 6 |
| Bract ratio | BM | -39.96 | 83.92 | 39.96 | 2 | 84.31 | 0.53 | 7 |

| Trait | Mode l | LogLikelihoo d | AIC | fOpt | k | AICc | Weight | Run |
|---|---|---|---|---|---|---|---|---|
| Bract ratio | EB | -39.96 | 85.92 | 39.96 | 3 | 86.72 | 0.16 | 7 |
| Bract ratio | OU | -39.96 | 85.92 | 39.96 | 3 | 86.72 | 0.16 | 7 |
| Bract ratio | VG | -39.96 | 85.92 | 39.96 | 3 | 86.72 | 0.16 | 7 |
| Bract ratio | BM | -39.96 | 83.92 | 39.96 | 2 | 84.31 | 0.52 | 8 |
| Bract ratio | EB | -39.91 | 85.83 | 39.91 | 3 | 86.63 | 0.16 | 8 |
| Bract ratio | NIG | -39.96 | 85.92 | 39.96 | 3 | 86.72 | 0.16 | 8 |
| Bract ratio | OU | -39.96 | 85.92 | 39.96 | 3 | 86.72 | 0.16 | 8 |
| Bract ratio | BM | -39.96 | 83.92 | 39.96 | 2 | 84.31 | 0.52 | 9 |
| Bract ratio | EB | -39.91 | 85.83 | 39.91 | 3 | 86.63 | 0.16 | 9 |
| Bract ratio | OU | -39.96 | 85.92 | 39.96 | 3 | 86.72 | 0.16 | 9 |
| Bract ratio | VG | -39.96 | 85.92 | 39.96 | 3 | 86.72 | 0.16 | 9 |
| Bract ratio | BM | -39.96 | 83.92 | 39.96 | 2 | 84.31 | 0.52 | 10 |
| Bract ratio | EB | -39.91 | 85.83 | 39.91 | 3 | 86.63 | 0.16 | 10 |
| Bract ratio | VG | -39.96 | 85.92 | 39.96 | 3 | 86.72 | 0.16 | 10 |
| Bract ratio | OU | -39.96 | 85.92 | 39.96 | 3 | 86.72 | 0.16 | 10 |
| Sepal ratio | BM | -51.78 | 107.56 | 51.78 | 2 | 107.95 | 0.46 | 1 |
| Sepal ratio | JN | -51.13 | 108.26 | 51.13 | 3 | 109.06 | 0.26 | 1 |
| Sepal ratio | OU | -51.77 | 109.54 | 51.77 | 3 | 110.34 | 0.14 | 1 |

| Trait | Model | LogLikelihood | AIC | fOpt | k | AICc | Weight | Run |
|---|---|---|---|---|---|---|---|---|
| Sepal ratio | EB | -51.78 | 109.56 | 51.78 | 3 | 110.36 | 0.14 | 1 |
| Sepal ratio | BM | -51.78 | 107.56 | 51.78 | 2 | 107.95 | 0.51 | 2 |
| Sepal ratio | VG | -51.64 | 109.28 | 51.64 | 3 | 110.08 | 0.18 | 2 |
| Sepal ratio | OU | -51.77 | 109.54 | 51.77 | 3 | 110.34 | 0.16 | 2 |
| Sepal ratio | EB | -51.78 | 109.56 | 51.78 | 3 | 110.36 | 0.15 | 2 |
| Sepal ratio | BM | -51.78 | 107.56 | 51.78 | 2 | 107.95 | 0.62 | 3 |
| Sepal ratio | OU | -51.77 | 109.54 | 51.77 | 3 | 110.34 | 0.19 | 3 |
| Sepal ratio | EB | -51.78 | 109.56 | 51.78 | 3 | 110.36 | 0.18 | 3 |
| Sepal ratio | JN | -54.52 | 115.05 | 54.52 | 3 | 115.85 | 0.01 | 3 |
| Sepal ratio | BM | -51.78 | 107.56 | 51.78 | 2 | 107.95 | 0.52 | 4 |
| Sepal ratio | JN | -51.72 | 109.44 | 51.72 | 3 | 110.24 | 0.17 | 4 |
| Sepal ratio | OU | -51.78 | 109.56 | 51.78 | 3 | 110.36 | 0.16 | 4 |
| Sepal ratio | EB | -51.78 | 109.56 | 51.78 | 3 | 110.36 | 0.16 | 4 |
| Sepal ratio | BM | -51.78 | 107.56 | 51.78 | 2 | 107.95 | 0.56 | 5 |

| Trait | Model | LogLikelihood | AIC | fOpt | k | AICc | Weight | Run |
|---|---|---|---|---|---|---|---|---|
| Sepal ratio | OU | -51.78 | 109.56 | 51.78 | 3 | 110.36 | 0.17 | 5 |
| Sepal ratio | EB | -51.78 | 109.56 | 51.78 | 3 | 110.36 | 0.17 | 5 |
| Sepal ratio | JN | -52.19 | 110.38 | 52.19 | 3 | 111.18 | 0.11 | 5 |
| Sepal ratio | BM | -51.78 | 107.56 | 51.78 | 2 | 107.95 | 0.38 | 6 |
| Sepal ratio | JN | -50.58 | 107.15 | 50.58 | 3 | 107.95 | 0.38 | 6 |
| Sepal ratio | OU | -51.78 | 109.56 | 51.78 | 3 | 110.36 | 0.12 | 6 |
| Sepal ratio | EB | -51.78 | 109.56 | 51.78 | 3 | 110.36 | 0.12 | 6 |
| Sepal ratio | BM | -51.78 | 107.56 | 51.78 | 2 | 107.95 | 0.50 | 7 |
| Sepal ratio | JN | -51.55 | 109.09 | 51.55 | 3 | 109.89 | 0.19 | 7 |
| Sepal ratio | OU | -51.77 | 109.54 | 51.77 | 3 | 110.34 | 0.15 | 7 |
| Sepal ratio | EB | -51.78 | 109.56 | 51.78 | 3 | 110.36 | 0.15 | 7 |
| Sepal ratio | BM | -51.78 | 107.56 | 51.78 | 2 | 107.95 | 0.51 | 8 |
| Sepal ratio | VG | -51.64 | 109.28 | 51.64 | 3 | 110.08 | 0.18 | 8 |
| Sepal ratio | OU | -51.78 | 109.56 | 51.78 | 3 | 110.36 | 0.15 | 8 |

| Trait | Model | LogLikelihood | AIC | fOpt | k | AICc | Weight | Run |
|---|---|---|---|---|---|---|---|---|
| Sepal ratio | EB | -51.78 | 109.56 | 51.78 | 3 | 110.36 | 0.15 | 8 |
| Sepal ratio | BM | -51.78 | 107.56 | 51.78 | 2 | 107.95 | 0.45 | 9 |
| Sepal ratio | JN | -51.03 | 108.07 | 51.03 | 3 | 108.87 | 0.28 | 9 |
| Sepal ratio | OU | -51.77 | 109.54 | 51.77 | 3 | 110.34 | 0.14 | 9 |
| Sepal ratio | EB | -51.78 | 109.56 | 51.78 | 3 | 110.36 | 0.13 | 9 |
| Sepal ratio | BM | -51.78 | 107.56 | 51.78 | 2 | 107.95 | 0.51 | 10 |
| Sepal ratio | JN | -51.55 | 109.09 | 51.55 | 3 | 109.89 | 0.19 | 10 |
| Sepal ratio | OU | -51.78 | 109.56 | 51.78 | 3 | 110.36 | 0.15 | 10 |
| Sepal ratio | EB | -51.78 | 109.56 | 51.78 | 3 | 110.36 | 0.15 | 10 |

**Table S3.** Model comparison results of the BioGeoBEARS analysis.

| Model | LnL | Number of parameters | Dispersal | Extinction | Jump speciation | AICc |
|---|---|---|---|---|---|---|
| DEC+J | -52.21 | 3 | 0.005 | $1 \times 10^{-12}$ | 0.019 | 111.2 |
| DIVALIKE+J | -55.41 | 3 | 0.01 | $1 \times 10^{-12}$ | 0.018 | 117.6 |
| BAYAREALIKE+J | -56.89 | 3 | 0.008 | $1 \times 10^{-12}$ | 0.02 | 120.6 |
| DIVALIKE | -61.35 | 2 | 0.022 | $1 \times 10^{-7}$ | 0 | 127.1 |
| DEC | -63.05 | 2 | 0.018 | 0.005 | 0 | 130.5 |
| BAYAREALIKE | -78.91 | 2 | 0.022 | 0.17 | 0 | 162.2 |

**Dataset S1 (separate file).** Morphological measurements and geographic occurrence data for species of the Septentrional Clade of Puya.
